# Catch-bond engineering surpasses high-affinity maturation for T cell receptor therapies against solid tumors

**DOI:** 10.64898/2026.07.26.740839

**Authors:** Tianqi Huang, Runyu Wang, Dinglin Zhang, Siqi Wu, Yuanhao Wang, Jianglai Wang, Zhengxu Ren, Sifan Wang, Lan Li, Yan Zhou, Xiaojing Wang, Yanling Bao, Mingyu Fan, Luxue Zhang, Junshuang Liu, Wenjie Yuan, Huairui Yuan, Sirui Li, Bo Sun, Fei Shao, Chenqi Xu, Guohui Li, Anhui Wang, Wenmao Huang, Xiang Zhao

## Abstract

The sensitivity of T cell receptors (TCRs) has traditionally been attributed to their affinity, a principle that has long underpinned both T cell biology and TCR engineering. Currently, high-affinity maturation remains the predominant strategy employed to enhance TCR sensitivity; however, this approach has been associated with severe off-target toxicities in clinical settings. In this study, using the only FDA-approved TCR-T therapy as a clinical reference, we demonstrate that force-induced catch bonds, rather than static affinity, primarily determine TCR sensitivity and facilitate the development of more effective TCR engineering strategies. Among polar and charged amino acids, histidine was determined to be the most effective residue for pinpointing engineering hotspots. TCRs engineered to form catch bonds exhibited superior performance compared to the FDA-approved TCR subjected to high-affinity maturation, improving the efficacy of TCR-T cell therapies against solid tumors without eliciting off-target toxicity or alloreactivity. Mechanistically, a correlation was observed between TCR sensitivity and the strength of catch bonds, whereas no such relationship was found with affinity. The duration of T cell–tumor cell interactions, immunological synapse formation, and the intensity of subsequent signaling are governed by catch bonds rather than by affinity. Structural and computational investigations have demonstrated that force-induced reconfiguration of the ligand-receptor interface, along with the formation of a novel hydrogen bond network, augments the specificity of interactions between the TCR and antigenic peptides. Furthermore, bispecific T cell engagers utilizing the TCR scaffold also formed catch bonds with tumor antigens, indicating intrinsic mechanosensory properties of the extracellular domains of TCRs and suggesting a novel therapeutic modality based on catch-bond-engineered TCRs.

## Introduction

The interaction between the T cell receptor (TCR) and the peptide-major histocompatibility complex (pMHC) is fundamental to T cell-mediated immunity. However, the mechanisms by which TCRs discriminate between antigenic peptides differing by as little as a single amino acid remain incompletely understood. Traditionally, static affinity has been regarded as the principal determinant of TCR sensitivity, constituting a central paradigm in TCR biology^1,2^. Parameters such as three-dimensional (3D) affinity, 3D on-rate, 3D off-rate, two-dimensional (2D) affinity, and many others have all been proposed to correlate closely with TCR sensitivity^3–11^. Nonetheless, conflicting reports have emerged, resulting in ambiguity within the field.

Recent studies have revealed that TCRs function as mechanosensors and can form catch bonds with pMHC molecules^12–36^. Catch bonds are characterized by an increase in the lifetime of the receptor-ligand interaction under mechanical shear force. The 2D off-rate of TCRs under applied force serves as a quantitative measure of catch bond formation. This phenomenon may be more critical than static affinity, as high-affinity TCRs may fail to be activated by pMHC due to slip bond formation rather than catch bond formation^33,36^. Moreover, the strength of TCR signaling has been suggested to correlate with the extent of catch bond formation^33,34^.

TCR-engineered T cell therapies exploit the specificity of the TCR-pMHC interaction to equip patient T cells with exogenous TCRs targeting tumor cells. The isolation of potent TCRs is challenging due to central tolerance mechanisms against many tumor-associated antigens derived from self-proteins. Consequently, protein engineering of TCRs is often necessary to enhance therapeutic efficacy. Over the past three decades, affinity maturation—analogous to approaches used in antibody engineering—has been the predominant strategy for TCR optimization. Notably, both the only FDA-approved TCR-T cell therapy and the sole approved TCR-based T cell engager (TCE) therapy utilize high-affinity-matured TCR variants^37–45^. However, affinity maturation has limitations, including the risk of severe off-target toxicities observed clinically and the fact that the highest affinity variants are not necessarily the most sensitive^46–48^. Engineering catch bonds between TCR and pMHC represents an alternative strategy to augment TCR sensitivity^33–35^. Yet, it remains uncertain whether catch bond engineering consistently offers advantages over affinity maturation across diverse TCR engineering applications.

In the present study, using the only FDA-approved high-affinity-matured TCR-T cell therapy as a clinical benchmark, we demonstrate that catch bond formation, rather than affinity, governs TCR sensitivity. Among charged and polar amino acids, histidine was identified as the most effective residue for identifying engineering hotspots on TCRs. Through histidine mutagenesis and combinatorial engineering of hotspots identified from histidine scanning, catch bond-engineered TCR variants exhibited superior performance compared to high-affinity-matured TCRs, showing enhanced in vivo efficacy without associated toxicity. Mechanistically, catch bond formation, rather than affinity, appears to drive the evolution of TCR sensitivity, as catch bond strength—but not affinity—correlates with TCR signaling potency. Enhanced catch bond strength facilitates prolonged interactions between tumor cells and T cells, promotes the formation of immunological synapse, and results in elevated phosphorylation levels of molecules involved in TCR downstream signaling pathways. Catch bonds enable the formation of force-induced hydrogen bonds, contribute to the reorganization of the TCR-pMHC interface, and enhance specific interactions with peptides. Furthermore, bispecific T cell engagers (BiTEs) constructed on a TCR scaffold also demonstrated catch bond formation with pMHC, underscoring the intrinsic mechanosensory properties of TCRs. Notably, catch bond-engineered BiTEs delayed tumor progression, whereas high-affinity-matured BiTEs failed to control tumor growth, suggesting that BiTEs may serve as an alternative platform for catch bond-based TCR therapeutics.

## Results

### Histidine plays a predominant role in the formation of new catch bonds

The FDA-approved TCR-T cell therapy, Tecelra, was selected as the clinical research benchmark (Fig. 1A). The complementarity-determining region (CDR) sequences of the wild-type (WT) TCR are presented in Fig. 1B. The TCR variant employed in the TCR-T cell therapy is the high-affinity matured TCR4 variant, whose CDR sequences are also detailed in Fig. 1B. Relative to the WT TCR, TCR4 harbors two mutations located in CDR2α and CDR3β (Fig. 1B) and exhibits an approximately 1000-fold increase in potency (Fig. 1C-D). To explore whether catch bond-driven evolution could yield a high-potency, low-affinity TCR variant surpassing the therapeutic efficacy of the high-affinity matured TCR4 *in vivo*, we undertook catch bond engineering of the WT TCR. Given that hydrogen bonds and salt bridges contribute to catch bond formation, we conducted a comprehensive amino acid scanning of the CDRs using all polar and charged residues to identify the most effective amino acids for establishing catch bonds (Fig. S1-S3). The scanning revealed distinct hotspot positions for different amino acids, indicating that certain hotspots may employ unique mechanisms influenced by the chemical properties of amino acid side chains.

**Fig. 1.**
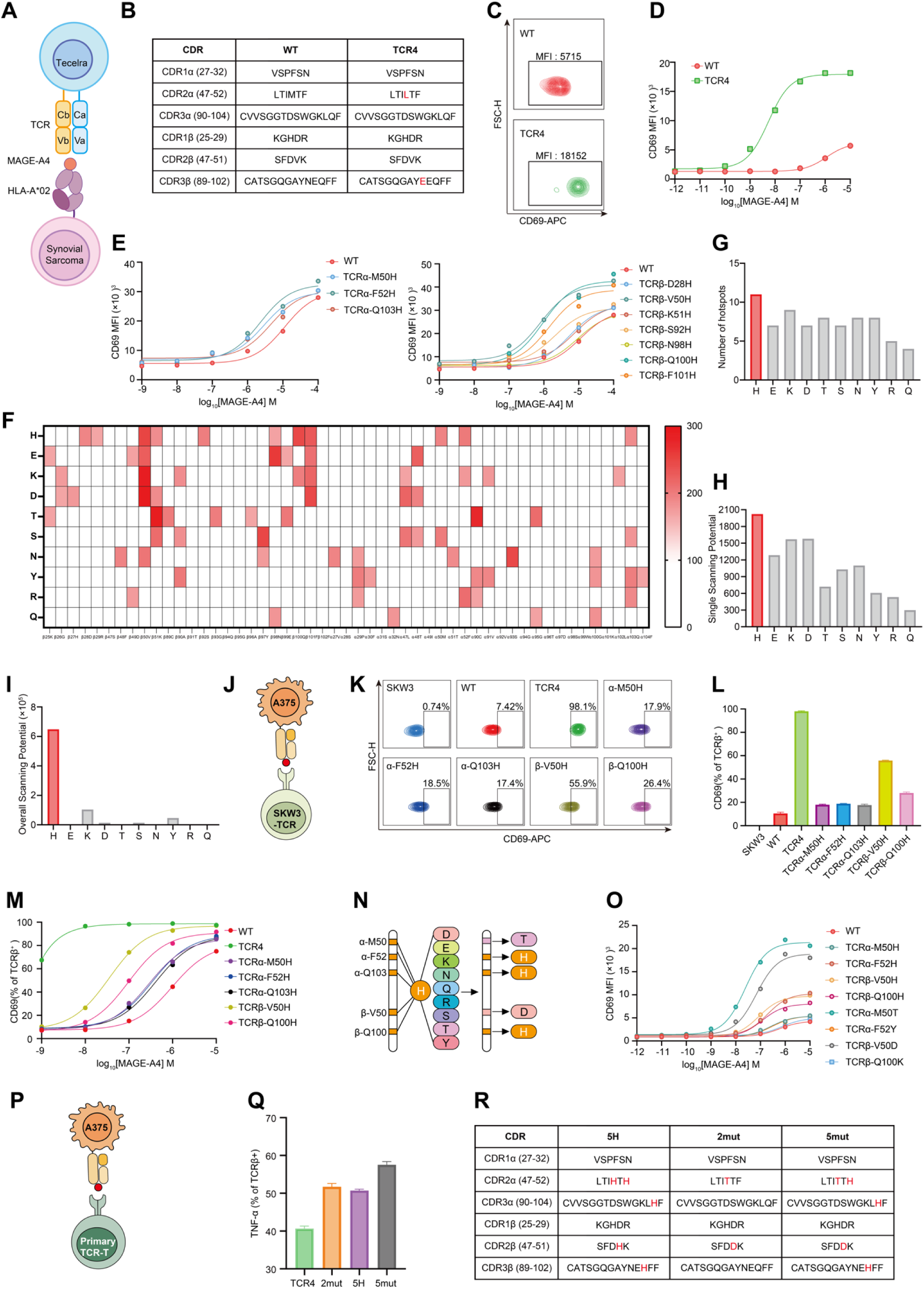
Catch bond engineering of the WT TCR. (A) Schematic of TCR4 (Tecelra)-transduced T cells killing HLA-A2-MAGE-A4-positive tumor cells. (B) CDRs sequence of the WT TCR and TCR4. (C) Sensitivity of the WT TCR and TCR4. WT TCR- or TCR4-transduced SKW3 cells were activated by T2 cells pulsed with 10^-5^ M MAGE-A4 peptides and CD69 upregulation in SKW3 cells were analyzed by flow cytometry. Representative flow plots were shown. (D) Sensitivity of the WT TCR and TCR4. WT TCR- or TCR4-transduced SKW3 cells were activated by T2 cells pulsed with titrated MAGE-A4 peptides and CD69 upregulation in SKW3 cells were analyzed by flow cytometry. (E) Hotspots of the WT TCR revealed by histidine scanning. Single-histidine-mutated TCR-transduced SKW3 cells were activated by T2 cells pulsed with titrated MAGE-A4 peptides. CD69 upregulation in SKW3 cells were analyzed by flow cytometry. Variants with higher CD69 upregulation compared to the WT TCR were shown. (F) Heatmap of polar and charged amino acid scanning of hotspots. Single-polar or charged amino acid-mutated TCR-transduced SKW3 cells were activated by T2 cells pulsed with titrated MAGE-A4 peptides. CD69 upregulation in SKW3 cells were analyzed by flow cytometry. The variants with anti-CD69 MFI higher than the anti-CD69 MFI of WT TCR were summarized in the heatmap showing the species of amino acids, the position of hotspots, and the Emax. All data were collected at a peptide concentration of 10⁻⁵ M. (G) Ranking of polar and charged amino acids in terms of the number of hotspots identified from the respective scanning. (H) The mutation expectation value for each residue position was defined based on the number and strength of activation enhancements induced by all possible substitutions to polar or charged amino acids at that position. For each kind of polar or charged amino acid, the single scanning potential was then calculated as the sum of the mutation expectation value across all hotspots, and polar and charged amino acids were ranked accordingly. (I) The activation probability value for each residue position was defined as the number of polar or charged amino acid substitutions that enhanced activation at that position. For each kind of polar or charged amino acid, the overall scanning potential was then calculated as the multiplication of the activation expectation value across all hotspots, and polar and charged amino acids were ranked accordingly. (J) Schematic of coculture of HLA-A2-MAGE-A4-positive A375 tumor cells with TCR-transduced SKW3 cells. (K) Sensitivity of top 5 potent single-histidine-mutated TCR variants. TCR-transduced SKW3 cells were cocultured with A375 cells. CD69 upregulation in SKW3 cells were analyzed by flow cytometry. Representative flow plots were shown. (L) Sensitivity of top 5 potent single-histidine-mutated TCR variants. TCR-transduced SKW3 cells were cocultured with A375 cells. CD69 upregulation in SKW3 cells were analyzed by flow cytometry. (M) Sensitivity of top 5 potent single-histidine-mutated TCR variants. TCR-transduced SKW3 cells were activated by T2 cells pulsed with titrated MAGE-A4 peptides and CD69 upregulation in SKW3 cells were analyzed by flow cytometry. (N) Schematic of combinatory engineering based on the histidine scanning hotspots. The most potent histidine scanning hotspots were further tested with mutagenesis into all the polar and charged amino acids. 2-5 hotspots may be mutated simultaneously. (O) Sensitivity of top 5 potent histidine hotspots mutated to histidine or certain amino acids which exhibited the most or the secondary potency at the respective hotspots. TCR-transduced SKW3 cells were activated by T2 cells pulsed with titrated MAGE-A4 peptides and CD69 upregulation in SKW3 cells were analyzed by flow cytometry. (P) Schematic of coculture of HLA-A2-MAGE-A4-positive A375 tumor cells with TCR-transduced primary T cells. (Q) Sensitivity of different combinatory engineered TCR variants. TCR-transduced primary T cells were cocultured with A375 cells. TNF expression in primary T cells were analyzed by flow cytometry. (R) CDRs sequence of the 2mut, 5H, and 5mut TCR variants. Data are representative of three experiments. Data are shown as means ± SD of technical duplicates. Statistical analysis was performed using two-tailed unpaired Student’s *t*-tests. n.s., not significant; \**P* < 0.05; \*\**P* < 0.01; \*\*\**P* < 0.001; \*\*\*\**P* < 0.0001.

Using TCR-transduced SKW3 cells, histidine scanning of the WT TCR identified ten hotspots where single histidine substitutions enhanced activation relative to the WT TCR (Fig. 1E). Among all polar and charged amino acids tested, histidine scanning yielded the greatest number of variants with improved activation and demonstrated the highest efficiency in pinpointing hotspots amenable to mutagenesis into multiple kinds of amino acids for activation enhancement (Fig. 1F-I). However, co-culture experiments involving TCR-transduced SKW3 cells and HLA-A2-MAGE-A4-positive A375 cells revealed that none of the single histidine-mutated variants outperformed TCR4 (Fig. 1J-L). Furthermore, TCR4 remained approximately 100-fold more potent than the most effective single histidine-mutated variant, TCRβ-V50H, in TCR-transduced SKW3 cells (Fig. 1M).

Subsequently, two engineering strategies were employed: first, several hotspots identified via histidine scanning were mutated to other polar or charged amino acids to determine which substitutions maximized TCR signaling strength; second, the top five hotspots from the histidine scanning were combined in pairs, triplets, quadruplets, or quintuplets of mutations (Fig. 1N). Some hotspots, such as TCRα-F52 and TCRβ-Q100, exhibited maximal potency with histidine substitutions, whereas others, including TCRα-M50T and TCRβ-V50D, achieved the highest signaling strength with alternative amino acid substitutions (Fig. 1O, Extended Data Fig. 4A-E). Using TCR-transduced primary T cells, three representative variants—2mut, 5H, and 5mut—were identified that surpassed TCR4 in terms of TCR activation and were selected for further analysis (Fig. 1P-R, Extended Data Fig. 4F-H, Table S1).

### Catch bond engineering outperforms high affinity maturation using the FDA-approved TCR as a clinical benchmark

Utilizing primary T cells transduced with TCRs, we observed that whereas the WT TCR exhibited co-receptor dependency, both engineered TCR variants and TCR4 functioned independently of co-receptors (Fig. 2A-E). In proliferation assays using TCR-transduced primary T cells stimulated with A375 tumor cells, engineered TCRs demonstrated enhanced proliferative responses relative to TCR4 and WT TCR. Although cell viability declined after day 8, the 5mut TCR variant maintained the highest T cell counts (Fig. 2F-G). Subsequent evaluation of repetitive cytotoxic activity against A375 tumor cells revealed that engineered TCR variants consistently mediated greater tumor cell killing compared to WT TCR. Notably, the 5mut TCR exhibited superior cytotoxicity in each round relative to the affinity-matured TCR4 (Fig. 2I-N). In the second and third rounds of killing assays, peripheral blood mononuclear cells (PBMCs) and WT TCR-T cells failed to suppress tumor proliferation, whereas engineered TCR-T cells effectively eliminated tumor cells, with the 5mut variant outperforming TCR4 across all rounds (Fig. 2M-N). Following three rounds of cytotoxic activity, engineered TCR variants displayed reduced expression of exhaustion markers PD-1 and TIM-3 compared to TCR4, suggesting that enhanced cytotoxicity was not accompanied by increased T cell exhaustion (Fig. 2O-S).

**Fig. 2.**
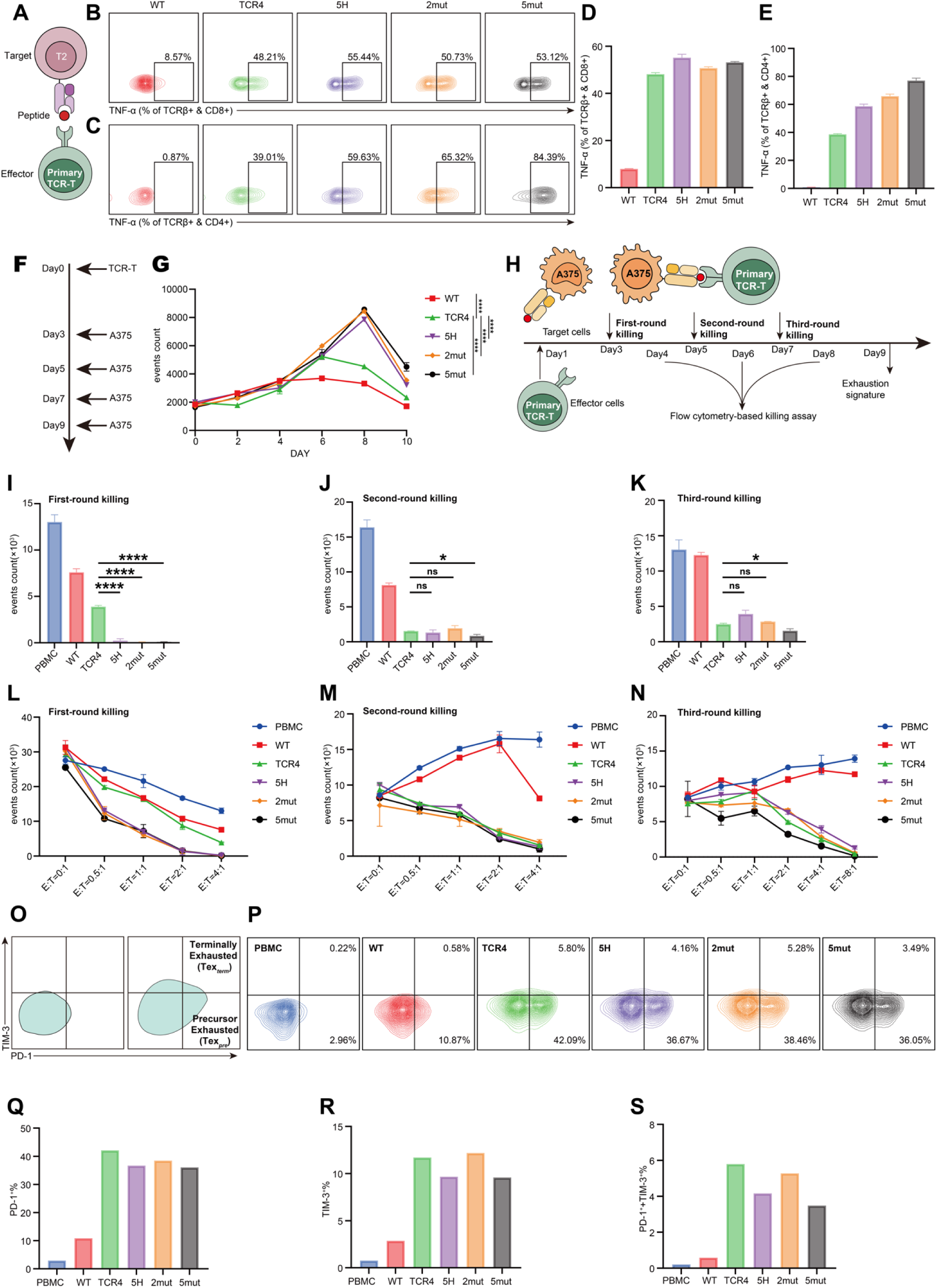
Catch-bond-engineered TCR-T cell therapy *in vitro*. (A) Schematic of coculture of primary TCR-T cells with T2 cells pulsed with MAGE-A4 peptides. (B) Co-receptor dependency of the selected TCR variants. TCR-transduced primary T cells were activated by T2 cells pulsed with 10^-5^ M MAGE-A4 peptides. T cells were stained with anti-TNF, anti-CD4, and anti-CD8. Representative flow plots of CD8^+^ T cells were shown. (C) Co-receptor dependency of the selected TCR variants. TCR-transduced primary T cells were activated by T2 cells pulsed with 10^-5^ M MAGE-A4 peptides. T cells were stained with anti-TNF, anti-CD4, and anti-CD8. Representative flow plots of CD4^+^ T cells were shown. (D) TNF production of the selected TCR variants in CD8^+^ T cells. TCR-transduced primary T cells were activated by T2 cells pulsed with 10^-5^ M MAGE-A4 peptides for 6 hours. T cells were stained with anti-TNF, anti-CD4, and anti-CD8. (E) TNF production of the selected TCR variants in CD4^+^ T cells. TCR-transduced primary T cells were activated by T2 cells pulsed with 10^-5^ M MAGE-A4 peptides for 6 hours. T cells were stained with anti-TNF, anti-CD4, and anti-CD8. (F) Schematic of antigen-induced proliferation of primary TCR-T cells. (G) Antigen-induced proliferation of TCR variants-transduced primary T cells. (H) Schematic of repetitive killing of A375 cells by primary TCR-T cells. (I) Cytotoxicity of the selected TCR variants in the first-round killing experiment. TCR variants-transduced primary T cells were cocultured with A375 cells at effectors: target (E: T) ratio of 4:1 for 24 hours. Leftover alive A375 cells were counted. (J) Cytotoxicity of the selected TCR variants in the second-round killing experiment. TCR variants-transduced primary T cells were cocultured with A375 cells at effectors: target (E: T) ratio of 4:1 for 24 hours. Leftover alive A375 cells were counted. (K) Cytotoxicity of the selected TCR variants in the third-round killing experiment. TCR variants-transduced primary T cells were cocultured with A375 cells at effectors: target (E: T) ratio of 4:1 for 24 hours. Leftover alive A375 cells were counted. (L) Cytotoxicity of the selected TCR variants in the first-round killing experiment. TCR variants-transduced primary T cells were cocultured with A375 cells at different effectors: target (E: T) ratios for 24 hours. Leftover alive A375 cells were counted. (M) Cytotoxicity of the selected TCR variants in the second-round killing experiment. TCR variants-transduced primary T cells were cocultured with A375 cells at different effectors: target (E: T) ratios for 24 hours. Leftover alive A375 cells were counted. (N) Cytotoxicity of the selected TCR variants in the third-round killing experiment. TCR variants-transduced primary T cells were cocultured with A375 cells at different effectors: target (E: T) ratios for 24 hours. Leftover alive A375 cells were counted. (O) Schematic of exhaustion phenotype of T cells based on PD-1 and TIM-3 gating. (P) Representative flow plots of PD-1 and TIM-3 staining of TCR variants-transduced primary T cells after three rounds of killing experiments. (Q) PD-1-positive percentage of TCR variants-transduced primary T cells after three rounds of killing experiments. (R) TIM-3-positive percentage of TCR variants-transduced primary T cells after three rounds of killing experiments. (S) PD-1 and TIM-3 double-positive percentage of TCR variants-transduced primary T cells after three rounds of killing experiments. Data are representative of three experiments. Data are shown as means ± SD of technical duplicates. Statistical analysis was performed using two-tailed unpaired Student’s *t*-tests. n.s., not significant; \**P* < 0.05; \*\**P* < 0.01; \*\*\**P* < 0.001; \*\*\*\**P* < 0.0001.

In an *in vivo* model, immunodeficient mice bearing A375 tumors received adoptive transfer of primary TCR-T cells (Fig. 3A). The 5mut TCR variant significantly delayed tumor progression relative to the affinity-matured TCR4 (Fig. 3B-E). Importantly, treatment with engineered TCR-T cells did not induce toxicity or weight loss (Fig. 3C). Moreover, mice treated with 5mut TCR-T cells exhibited significantly prolonged survival compared to those receiving TCR4 TCR-T cells (Fig. 3E). We did not observe significant advantages of treatment from 2mut TCR-T cells or 5H TCR-T cells compared to TCR4 TCR-T cells (Extended Data Fig. 4I-M).

**Fig. 3.**
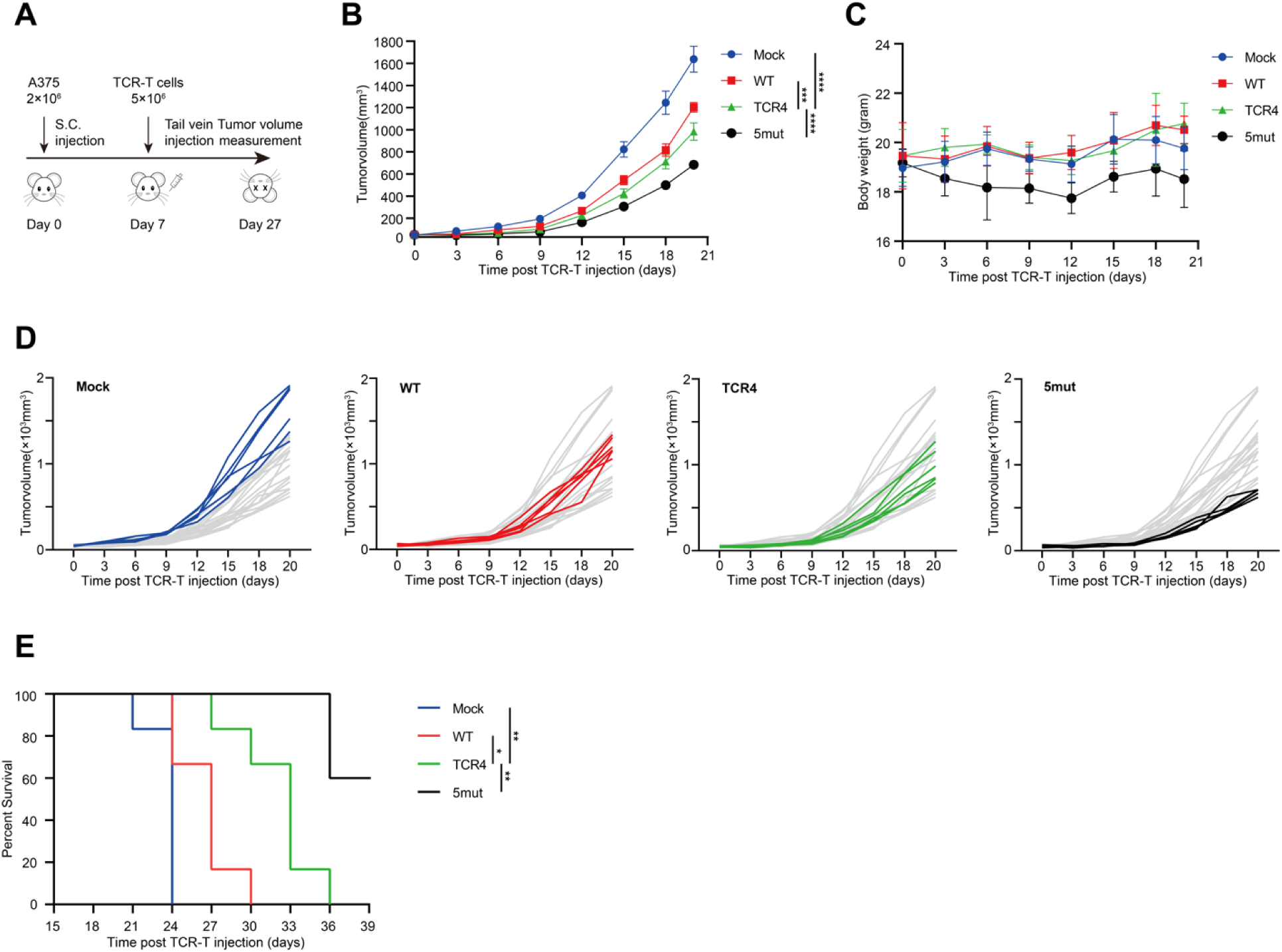
Catch-bond-engineered TCR-T cell therapy *in vivo*. (A) Schematic of testing primary TCR-T cell therapy in tumor-bearing immunodeficient mice *in vivo*. (B) Tumor volume of mice post TCR-T cells injection. Immunodeficient mice were implanted with A375 tumor cells followed by adoptive transfer of TCR variants-transduced primary T cells.(C) Body weight of mice post TCR-T cells injection. (D) Tumor volume of individual mouse post TCR-T cells injection (E) Survival curve of mice post TCR-T cells injection. Data are representative of two experiments. Data are shown as means ± SD of technical duplicates. Statistical analysis was performed using two-tailed unpaired Student’s *t*-tests. n.s., not significant; \**P* < 0.05; \*\**P* < 0.01; \*\*\**P* < 0.001; \*\*\*\**P* < 0.0001.

To assess specificity, we conducted an X-scan analysis for TCR4 and the engineered 5mut TCR variant (Fig. 4A-B). Both TCRs demonstrated similar tolerance patterns, with most amino acid substitutions at positions P5-P9 being permissible. Using a 10% tolerance threshold, we generated binding motifs to predict potential off-target peptides for each TCR. Due to the large number of predicted peptides, a position weight matrix derived from the X-scan heatmap was employed to rank candidates^49^. We synthesized the top 100 peptides (Table S2). Functional assays using TCR-transduced SKW3 cells revealed that both the TCR4 and the 5mut TCR variants exhibited no cross-reactivity (Fig. 4C-D). Furthermore, co-culture experiments with TCR-transduced SKW3 cells and panels of HLA-A2^+^ MAGE-A4^-^ or HLA-A2^-^ MAGE-A4^+^ tumor cells demonstrated no evidence of alloreactivity for the engineered TCR variants (Fig. 4E-H).

**Fig. 4.**
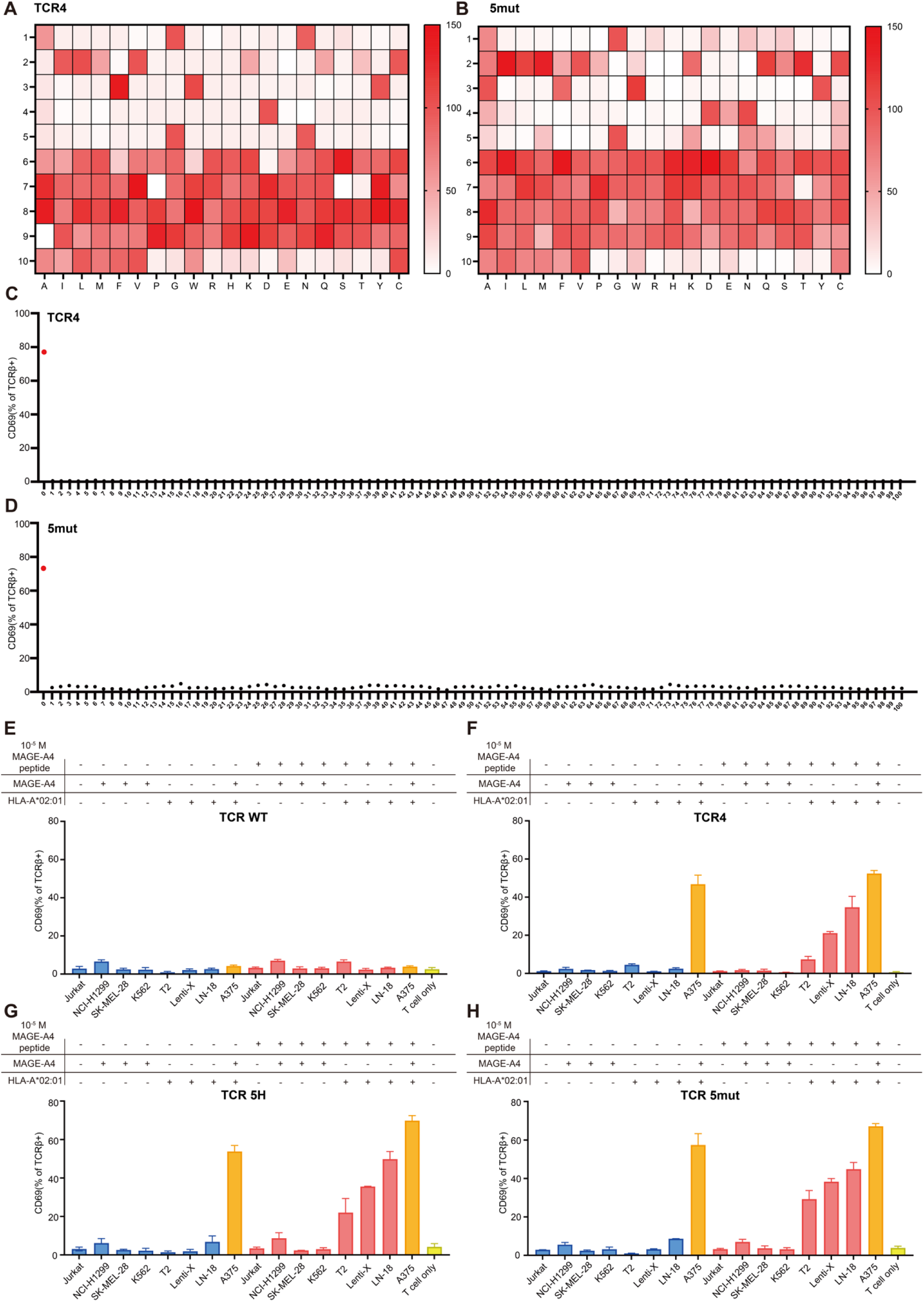
The off-target toxicity and alloreactivity of catch-bond-engineered TCR. (A) X-scan of the TCR4 variant. TCR4-transduced SKW3 cells were cocultured with T2 cells pulsed with MAGE-A4 peptide variants. SKW3 cells were stained with anti-CD69-APC. The MFI of TCR4-transduced SKW3 cells stimulated by MAGE-A4 peptide were set as the 100% level. (B) X-scan of the 5mut TCR variant. 5mut TCR-transduced SKW3 cells were cocultured with T2 cells pulsed with MAGE-A4 peptide variants. SKW3 cells were stained with anti-CD69-APC. The MFI of 5mut TCR-transduced SKW3 cells stimulated by MAGE-A4 peptide were set as the 100% level. (C) Off-target peptides screening for the TCR4. TCR4-transduced SKW3 cells were cocultured with T2 cells pulsed with different predicted off-target peptides. SKW3 cells were stained with anti-CD69-APC. (D) Off-target peptides screening for the 5mut TCR. 5mut TCR-transduced SKW3 cells were cocultured with T2 cells pulsed with different predicted off-target peptides. SKW3 cells were stained with anti-CD69-APC. (E) Alloreactivity and off-target toxicity of the WT TCR. WT TCR-transduced SKW3 cells were cocultured with panels of tumor cell lines. SKW3 cells were stained with anti-CD69-APC. (F) Alloreactivity and off-target toxicity of the TCR4 variant. TCR4-transduced SKW3 cells were cocultured with panels of tumor cell lines. SKW3 cells were stained with anti-CD69-APC. (G) Alloreactivity and off-target toxicity of the 5H TCR variant. 5H TCR-transduced SKW3 cells were cocultured with panels of tumor cell lines. SKW3 cells were stained with anti-CD69-APC. (H) Alloreactivity and off-target toxicity of the 5mut TCR variant. 5mut TCR-transduced SKW3 cells were cocultured with panels of tumor cell lines. SKW3 cells were stained with anti-CD69-APC. Data are representative of three experiments. Data are shown as means ± SD of technical duplicates. Statistical analysis was performed using two-tailed unpaired Student’s *t*-tests. n.s., not significant; \**P* < 0.05; \*\**P* < 0.01; \*\*\**P* < 0.001; \*\*\*\**P* < 0.0001.

### The regulatory mechanism underlying TCR sensitivity

We examined the underlying mechanism by which TCR sensitivity is determined (Fig. 5A). The 3D binding affinity between various TCR variants and the HLA-A2-MAGE-A4 complex was quantified using surface plasmon resonance. Notably, TCR4 exhibited an 11-fold higher affinity relative to the WT TCR. In contrast, all engineered TCR variants demonstrated substantially lower affinities compared to TCR4. The most efficacious variant, designated 5mut TCR, displayed an affinity of 173 nM, representing only a marginal increase relative to the WT TCR (Fig. 5B). These findings indicate that catch-bond engineering constitutes a strategy for generating TCRs characterized by low affinity yet high functional potency.

**Fig. 5.**
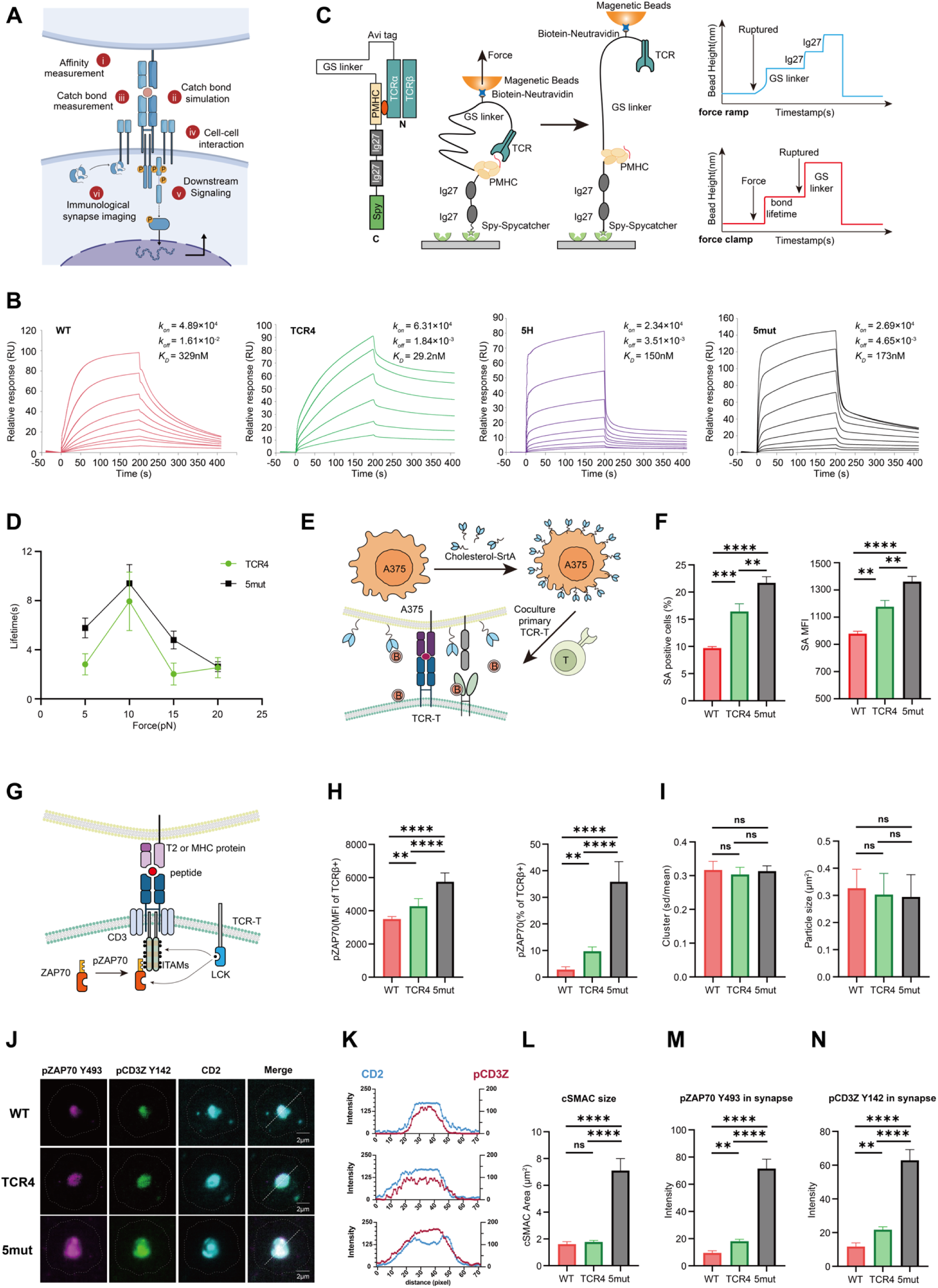
The mechanism of enhanced sensitivity of catch-bond-engineered TCRs. (A) Schematic of mechanism research of TCR sensitivity. (B) 3D binding affinity between different TCR variants and HLA-A2-MAGE-A4. The biotinylated HLA-A2-MAGE-A4 were immobilized on the streptavidin chip and different concentrations of recombinant extracellular TCR variants proteins were flowed through the channels of chips. (C) Schematic of principle of catch bond measurement by the magnetic tweezer. (D) Catch bond measurement between different TCR variants and HLA-A2-MAGE-A4 (E) Schematic of principle of cell-cell interaction strength measurement by the sortase assay. (F) Cell-cell interaction strength measurement between A375 cells and different TCR variants-transduced primary T cells by the engineered-sortase assay. (G) Schematic of ZAP70 phosphorylation level detection by flow cytometry. (H) pZAP70 level of different TCR variants-transduced Jurkat cells activated by T2 cells pulsed with MAGE-A4 peptides. (I) TCR clustering in the antigen-free state. Jurkat cells expressing the indicated TCRs were stained with anti-TCRβ antibody, and cluster size was quantified by SD/mean or cluster particle size. WT TCR, n = 38; TCR4, n = 41; mut5, n = 40. (J) Immunological synapse formation of WT, TCR4, and 5mut TCRs. Jurkat cells expressing the indicated TCRs and CD2-BFP were stimulated on supported lipid bilayers (SLBs) functionalized with pMHC, ICAM-1, CD58, ICOSL, and CD80 for 30 min. Representative images of immunological synapses formed by the indicated TCRs. Scale bar, 2 μm. (K) Overlay of phosphorylated TCR (pCD3ζ) and CD2 corolla. (L) Quantification of cSMAC size, defined by the area of pCD3ζ. (M) Quantification of ZAP70 Y493 phosphorylation at the immunological synapse. (N) Quantification of CD3ζ Y142 phosphorylation at the immunological synapse. WT TCR, n = 34; TCR4, n = 42; 5mut, n = 36 for (K)-(N). Data are representative of three experiments. Data are shown as means ± SD of technical duplicates. Statistical analysis was performed using two-tailed unpaired Student’s *t*-tests. n.s., not significant; \**P* < 0.05; \*\**P* < 0.01; \*\*\**P* < 0.001; \*\*\*\**P* < 0.0001.

To further elucidate the biophysical properties of TCR-pMHC interactions, we assessed catch bond formation between TCR variants and HLA-A2-MAGE-A4 using magnetic tweezers (Fig. 5C). To directly quantify the force-dependent interaction between TCR variants and HLA-A2-MAGE-A4 at the single-molecule level, we engineered recombinant protein constructs where TCR variants were fused to pMHC through an unstructured GS linker (Fig. 5C, left). For single-molecule measurements, individual protein constructs were tethered between a Neutravidin-coated superparamagnetic microbead and a SpyCatcher-functionalized glass surface via biotin–Neutravidin and SpyTag–SpyCatcher interactions (Fig. 5C, middle). Mechanical force was applied to the bead using magnetic tweezers, stretching the TCR-pMHC complex along the C-termini of both TCR and pMHC, mimicking the native force geometry during cell-cell contact. Once the TCR rupture from pMHC, the bead height would increase by an amount of step size equal to the extension change of the protein construct from the released GS linker (force ramp, Fig. 5C, right).

To directly determine the force-dependent stability of the TCR-pMHC interaction, we quantified bond lifetimes using a force-jump assay (force clamp, Fig. 5C, right). We measured the bond lifetime of TCR-pMHC complexes from ∼5 pN to ∼20 pN. Specifically, the lifetime of both complexes initially increased with increasing force, reached its maximum at ∼10 pN, and decreased at higher forces, exhibiting typical catch-bond behavior (Fig. 5D). Importantly, we observed that catch-bond-engineered TCR variants exhibited increased peak bond lifetimes relative to TCR4 (Fig. 5D).

To investigate the influence of TCR-pMHC interactions on cell-cell adhesion, we utilized a sortase assay. Cholesterol-conjugated Sortase-labeled A375 cells were co-cultured with primary TCR-transduced T cells, enabling the sortase-mediated covalent conjugation of a biotinylated substrate to the surface of T cells engaged in close contact with A375 cells. The extent of Streptavidin-647 staining on TCR-T cells served as a proxy for the strength of cell-cell interactions (Fig. 5E). Results demonstrated that 5mut TCR-T cells exhibited significantly enhanced interaction strength compared to TCR4, suggesting that catch bond engineering facilitates more robust cell-cell adhesion (Fig. 5F).

To investigate the activation speed of early phosphorylation signaling upon TCR-pMHC interaction, we measured the phosphorylated ZAP70 levels of TCR WT and two TCR variants after 15 min of antigen stimulation (Fig. 5G). Results demonstrated that 5mut TCR-Jurkat T cells exhibited significantly higher phosphorylation of ZAP70 than TCR4, indicating a more rapid or robust early signaling response (Fig. 5H).

Further analyses using TCR-transduced Jurkat cells showed that, in the absence of antigen, neither TCR4 nor the 5mut TCR exhibited excessive clustering, as evidenced by comparable TCR microcluster indices (SD/mean) and similar microcluster particle sizes (Fig. 5I). Upon TCR engagement, the 5mut TCR formed a more optimal immune synapse morphology, characterized by a larger synapse area and a CD2-looped configuration (Fig. 5J-K). In addition, phosphorylated CD3 accumulated more efficiently at the synapse, leading to enhanced downstream signaling (Fig. 5L-N).

Collectively, these data indicate that engineering TCR catch bonds promotes the formation of a mature immunological synapse without inducing undesired TCR self-clustering under antigen-free conditions.

### Molecular dynamics simulation of the TCR–pMHC binding mechanism and catch bond formation

We further investigated the mechanistic role of residue mutations in modulating binding affinity and promoting catch bond formation between TCR variants and HLA-A2-MAGE-A4 (hereafter pMHC). To this end, both WT and 5mut in complex with pMHC were constructed as model systems based on AlphaFold3 predictions (Extended Data Fig. 5A). As a control, TCR4 complex was also generated.

We first compared the binding footprint and contact contribution of individual CDR loops in binding to pMHC. Mutations within CDR2α, CDR3α, and CDR3β of TCR4 and 5mut induced noticeable conformational changes relative to the WT. However, the intermolecular contacts of the TCR4 and 5mut complexes remained highly similar, making it difficult to distinguish their functional differences based solely on static models (Extended Data Fig. 5B-C). Previous research has indicated that the static crystal structure of the TCR-pMHC complex is not helpful for analyzing the catch bond mechanism, and that molecular dynamics (MD) simulation is a better approach for observing the dynamic binding involved in catch bond formation^34,36,50^. Given that static structural predictions were insufficient to resolve these differences, we next turned to molecular dynamics simulations to capture the dynamic behavior of the complexes and uncover the mechanistic basis of their functional divergence.

To refine the AlphaFold-predicted structures and explore their conformational landscapes, three independent 500 ns relaxion MD simulations were performed for each complex, yielding stable conformational ensembles (Extended Data Fig. 5D). Based on the resulting trajectories, the binding free energies between TCR variants and pMHC were calculated. As shown in Fig. 6A and 6B, the binding free energy of 5mut with pMHC lies between that of WT and TCR4, in agreement with surface plasmon resonance (SPR) measurements. We further examined the structural effects of these mutations on the complexes. Both the TCR4 and 5mut complex systems exhibited a greater number of interfacial hydrogen bonds and salt bridges compared, resulting in stronger binding affinities (Table S3). In the WT system, a hydrogen bond between Q94β and Q155 is present, which may dissociate under shear force and be replaced by alternative interactions (Fig. 6C and Table S3). In the TCR4 system, the N98Eβ mutation introduces new hydrogen bonds, including E98β–Q155 and E98β–P6(R), thereby perturbing the local interaction network and inducing additional interactions such as Y97β–P4(D), Y97β–T163, and D97α–K66. These newly formed interactions contribute approximately −35 kcal/mol to the binding free energy, significantly enhancing TCR-pMHC binding (Fig. 6D and Table S3). In the 5mut system, the M50Tα and F52Hα mutations promote the formation of a new hydrogen bond F30α–E166. Meanwhile, the Q100Hβ mutation alters the local interaction network, leading to additional hydrogen bonds such as Y97β–P4(D), Y97β–T163, and Q94β–Q155. These interactions contribute approximately −25 kcal/mol to binding, resulting in an intermediate affinity between WT and TCR4 (Fig. 6E and Table S3). Notably, several of these interactions, including F30α–E166, Y97β–T163, and Q94β–Q155, are capable of transitioning into catch bonds under shear force.

**Fig. 6.**
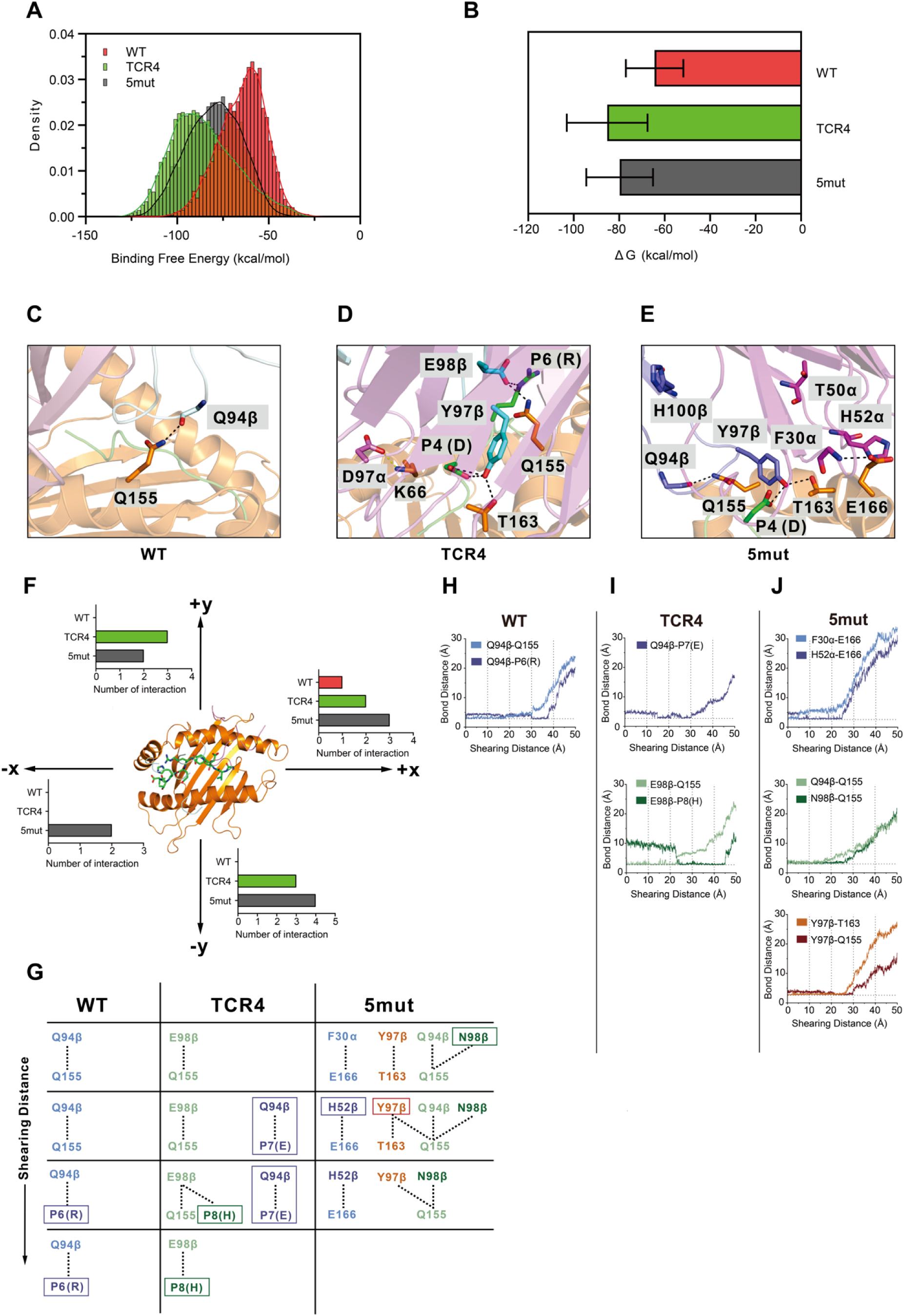
Dynamic Model of TCR-pMHC catch bond formation. (A) Distribution of binding free energies for WT TCR and its variants (TCR4 and 5mut) in complex with pMHC, calculated from three independent MD trajectories. (B) Statistical summary of binding free energies shown in (A). The 5mut variant exhibits an intermediate affinity between WT and TCR4, consistent with SPR measurements. (C) Close-up view of the interfacial interactions between WT and pMHC. TCRα-chain is shown in pink, β-chain in pale cyan, peptide in green, and HLA-A2 in orange. Key residues are displayed as sticks, with hydrogen bonds and salt bridges indicated by black dashed lines. The dominant interaction Q94β-Q155 is highlighted. (D) Same representation as in (C) for TCR4 with the TCRα-chain in violet and β-chain in cyan. Mutation N98Eβ remodels the interaction network, introducing additional contacts (e.g., E98β-Q155, E98β-P6(R), Y97β-P4(D), Y97β-T163, and D97α-K66), thereby strengthening binding affinity. (E) Same representation as in (C) for the 5mut variant with the TCRα-chain in magenta and β-chain in slate. Mutations (M50Tα, F52Hα, and Q100Hβ) induce new and redistributed interactions, including F30α-E166 and enhanced Y97β-mediated contacts, resulting in intermediate binding affinity. (F) Summary of newly formed interfacial interactions (primarily hydrogen bonds and salt bridges) during SMD simulations under shear in the ±x and ±y directions. TCR4 and 5mut exhibit more frequent formation of new interactions than WT, with 5mut showing the highest overall number. (G) Schematic representation of the dynamic evolution of interfacial interactions during forced dissociation along the +x direction for WT (left), TCR4 (middle), and 5mut (right). Newly formed interactions are highlighted in boxed regions and darker colors. Superscripts denote TCR residues, parentheses indicate peptide residues, and other residues belong to HLA-A2. (H) - (J) Bond distance versus shearing distance plots illustrating the dynamic behavior of representative interactions during dissociation under +x shear: (H) WT: transition from Q94β-Q155 to Q94β-P6(R); (I) TCR4: mutation-induced interactions (e.g., E98β-Q155) evolve into new contacts such as E98β-P8(H); (J) 5mut: multiple sequential interaction transitions (e.g., H52α-E166 to F30α-E166, Y97β-T163 to Y97β-Q155, Q94β-Q155 to N98β-Q155), supporting sustained resistance to dissociation and stronger catch bond behavior. Horizontal gray dashed lines show the equilibrium distance for the bond.

To further probe the dynamic evolution of molecular interactions during TCR-pMHC dissociation, we performed steered molecular dynamics (SMD) simulations. In these simulations, the center of mass of the TCR Vαβ domain was subjected to shear in four distinct directions relative to the pMHC (Fig. 6F and Methods), and the formation of transient interactions, including hydrogen bonds and salt bridges, was systematically quantified. Under shear in the ±x and ±y directions, both TCR4 and 5mut exhibited a higher number of newly formed interactions compared to WT (Fig. 6F, 6H-6J and Extended Data Fig. 6), with 5mut showing the greatest increase. To obtain atomic-level insights, we focused on dissociation under shear in the +x direction, where all three systems formed new interactions during the shearing process (Fig. 6G, Table S4, and Table S5). In the WT system, as the shearing distance increased, the original interaction Q94β–Q155 was replaced by a newly formed interaction Q94β–P6(R) (Fig. 6G and 6H, Extended Data Fig. 6A, and Movie S1). In the TCR4 system, following introduction of the N98Eβ mutation, a new interaction Q94β–P7(E) emerged during the early stage of shearing. Subsequently, the mutation-induced hydrogen bond E98β–Q155 transitioned into a new pair of interactions, E98β–P8(H), under shear force. This process effectively compensates for the loss of original interactions and reinforces the catch bond strength (Fig. 6G and 6I, Extended Data Fig. 6A, and Movie S2). In the 5mut system, multiple mutation-induced hydrogen bonds, including F30α–E166, Y97β–T163, and Q94β–Q155, were sequentially replaced by new interactions during the shearing process, such as H52α–E166, Y97β–Q155, and N98β–Q155. These interactions continuously resist TCR–pMHC dissociation, resulting in the strongest observed catch bond behavior (Fig. 6G and 6J, Extended Data Fig. 6A, and Movie S3).

Overall, these MD simulations provide mechanistic insights into how mutations in distinct regions regulate TCR-pMHC binding affinity by modulating the number of interfacial hydrogen bonds, and further tune catch bond strength by controlling the formation of new interactions under shear. Importantly, not all pre-existing interfacial interactions are converted into new interactions during shearing, which explains why the variant with the strongest binding affinity does not necessarily exhibit the strongest catch bond behavior.

### T cell engagers establish catch bonds with antigenic pMHC

BiTE molecules based on TCR scaffolds represent a significant alternative format for TCR-targeted therapeutics. Notably, the only FDA-approved TCR-BiTE utilizes a high-affinity matured TCR. However, it remained unclear whether BiTEs are capable of forming catch bonds with pMHC antigens. To address this, we designed and synthesized BiTE molecules and evaluated their effector functions (Fig. 7A). Using primary T cells, we observed that BiTEs engineered to enhance catch bond formation elicited significantly higher cytokine production (Fig. 7B–E) and cytotoxic activity (Fig. 7F–G) compared to the TCR4-BiTE. In an *in vivo* model, immunodeficient mice bearing A375 tumors were treated with adoptively transferred peripheral blood mononuclear cells (PBMCs) and intraperitoneal injections of BiTEs. The catch bond-engineered BiTE demonstrated a greater capacity to delay tumor progression relative to TCR4-BiTE (Fig. 7H–J). In contrast, the high-affinity matured TCR4-BiTE failed to control tumor growth when compared to the negative control (Fig. 7I). Catch bond formation was quantified using optical tweezers (Fig. 7K). Employing PBMCs, BiTE proteins, and pMHC complexes, we confirmed that BiTE molecules can form catch bonds with pMHC in the presence of CD3-expressing PBMC cells (Fig. 7L). Moreover, the catch bond-engineered BiTE exhibited a longer bond lifetime than the TCR4-BiTE (Fig. 7L). Collectively, these findings indicate that BiTEs engineered to enhance catch bond interactions outperform high-affinity matured BiTEs in tumor inhibition through a catch bond-mediated mechanism.

**Fig. 7.**
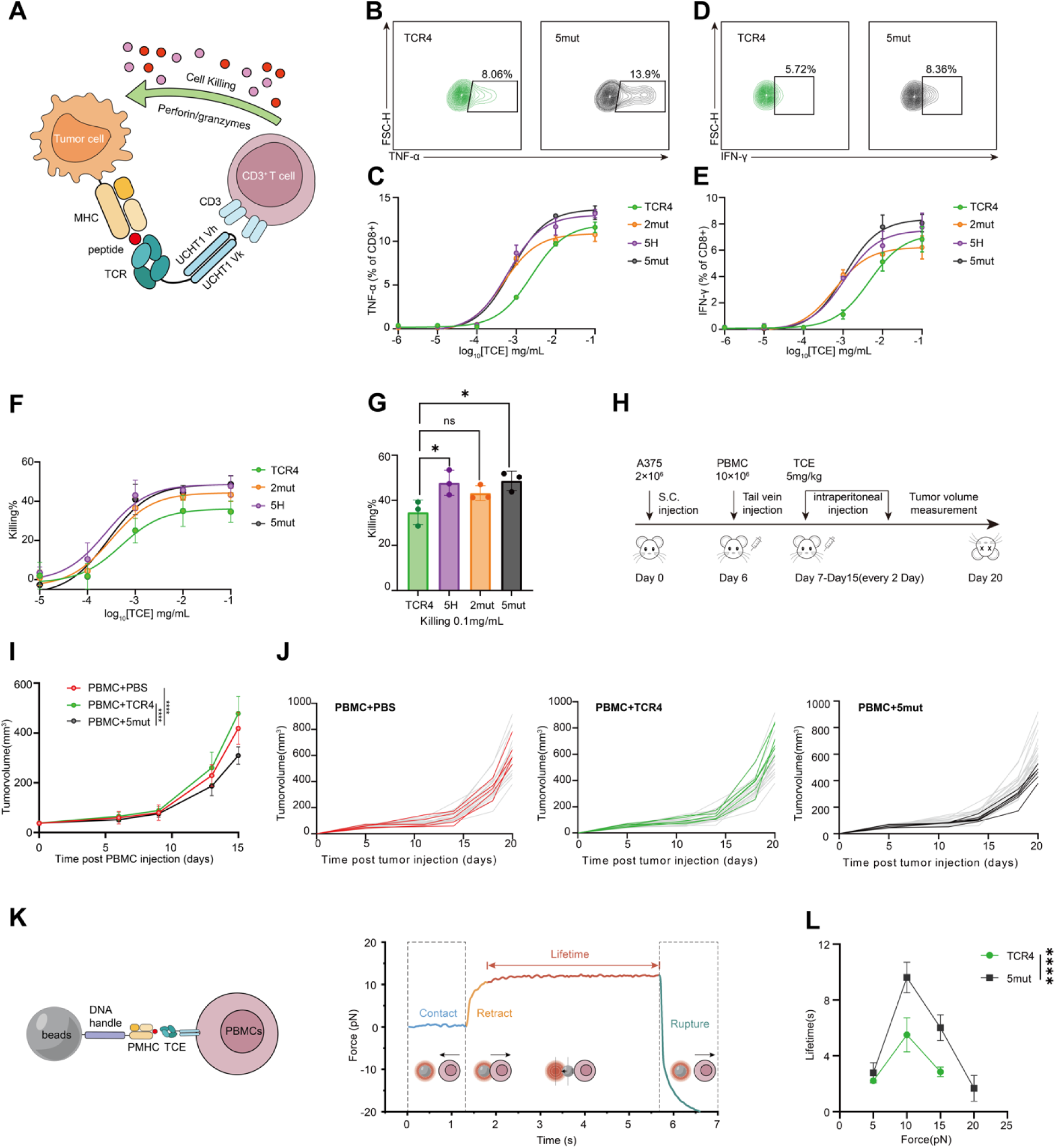
Efficacy of catch-bond-engineered T cell engagers *in vivo*. (A) Schematic of BiTE therapy. (B) TNF production of primary T cells activated by A375 cells and different BiTE. Representative flow plots of BiTE treatment at the concentration of 0.1 mg/mL were shown. (C) TNF production of primary T cells activated by A375 cells and different BiTE with titrated concentration. (D) IFN-γ production of primary T cells activated by A375 cells and different BiTE. Representative flow plots of BiTE treatment at the concentration of 0.1 mg/mL were shown. (E) IFN-γ production of primary T cells activated by A375 cells and different BiTE with titrated concentration. (F) Cytotoxicity of primary T cells induced by BiTE. Primary T cells were cocultured with A375 cells in the presence of different BiTE with titrated concentrations. (G) Cytotoxicity of primary T cells induced by BiTE. Primary T cells were cocultured with A375 cells in the presence of different BiTE at the concentration of 0.1 mg/mL. (H) Schematic of testing BiTE therapy in tumor-bearing immunodeficient mice *in vivo*. (I) Tumor volume of mice post PBMC injection. (J) Tumor volume of individual mouse post PBMC injection. (K) Schematic of catch bond measurement by optical tweezer. (L) Bond lifetime measurement between HLA-A2-MAGE-A4 and BiTE at 10 pN in the presence of PBMC cells. Data are representative of three experiments. Data are shown as means ± SD of technical duplicates. Statistical analysis was performed using two-tailed unpaired Student’s *t*-tests. n.s., not significant; \**P* < 0.05; \*\**P* < 0.01; \*\*\**P* < 0.001; \*\*\*\**P* < 0.0001.

## Discussion

The prevailing theory of T cell receptor (TCR) activation asserts that TCR sensitivity is chiefly determined by its inherent static affinity. Historically, TCR engineering has been largely guided by the affinity maturation paradigm, predicated on the intuitive notion that increased binding strength correlates with enhanced activation. In this study, through a direct comparative analysis between catch-bond-engineered TCRs and an FDA-approved affinity-matured TCR, we provide evidence that TCR sensitivity is regulated by force-induced catch bonds rather than static affinity alone. The approach of catch bond engineering signifies a conceptual shift, emphasizing dynamic, force-dependent interaction kinetics over static three-dimensional affinity. This strategy results in superior T cell activation and improved tumor eradication compared to high-affinity maturation, without compromising safety. Our results not only establish catch bond engineering as a more efficacious strategy than affinity maturation but also demonstrate its broad applicability across various therapeutic modalities, including TCR-engineered T cells and bispecific T cell engagers (BiTEs).

### Catch bonds govern TCR sensitivity more significantly than affinity

High binding affinity alone may not necessarily lead to robust TCR activation. Our findings offer a mechanistic insight into this phenomenon: TCR efficacy is strongly associated with the peak bond lifetime, a dynamic parameter reflecting the quality of catch bonds, rather than with equilibrium binding affinity. Although TCR4 demonstrated an 11-fold increase in affinity compared to the wild-type TCR, variants engineered to enhance catch bond formation exhibited lower affinity than TCR4 but showed significantly prolonged peak bond lifetimes and superior T cell activation in both *in vitro* and *in vivo* settings. These results underscore that optimizing force-dependent bond stabilization, rather than merely reducing the dissociation rate in the absence of force, is critical for effective TCR triggering.

We hypothesize that the formation of catch bonds necessitates precise geometric and chemical complementarity under mechanical load, functioning as a “mechano-checkpoint” that prevents recognition of peptides or HLA variants incapable of supporting the force-dependent reorganization of the interaction network. Consequently, catch bond engineering and the underlying catch bond mechanism finely regulate the interactions of specific residues under mechanical force, providing a force-dependent modality for antigen specificity. This intrinsic specificity may account for the absence of alloreactivity or off-target toxicity observed in our study.

### Histidine promotes the formation of catch bonds with the highest efficiency

A comprehensive evaluation of single-amino-acid scanning across all charged and polar residues demonstrated that histidine consistently identified the greatest number of hotspots associated with enhanced TCR sensitivity. The distinctive chemical characteristics of histidine—including its moderately sized side chain, capacity to function as both a hydrogen bond donor and acceptor, and its adjustable protonation state—are likely responsible for its ability to establish novel, force-sensitive interactions without inducing significant steric hindrance. We propose that histidine mutagenesis exemplifies a mechanism of “force-induced fitting” wherein the side chains of specific amino acids at particular positions adapt to facilitate catch bond formation under mechanical stress. Consequently, histidine scanning emerges as a versatile and effective initial strategy for engineering catch bonds, especially in contexts lacking detailed structural data. Although other charged and polar residues also revealed functional hotspots, their efficacy was variable, indicating that future engineering endeavors may benefit from integrating insights derived from multiple scanning methodologies.

### Engineering of catch bonds in TCRs preserves functional T cell states and enhances antitumor efficacy

Beyond immediate activation, TCRs engineered with catch bonds demonstrated superior antitumor activity and favorable differentiation profiles compared to TCRs matured for high affinity. In vivo studies revealed that TCR-T cells expressing catch-bond variants achieved improved tumor control. Importantly, although the TCR4 variant induced increased expression of exhaustion markers such as PD-1 and TIM-3, catch-bond variants maintained exhaustion profiles lower than those of TCR4. This observation supports the hypothesis that mechanical sensing mediated by catch bonds provides a more physiological activation signal, thereby promoting sustained T cell functionality without accelerating dysfunction. The enhanced repetitive cytotoxicity and proliferative capacity observed further underscore the potential of catch bond engineering to elicit more durable therapeutic responses. Consequently, catch bond engineering may optimize not only the magnitude but also the quality of TCR signaling.

### Extension of catch bond engineering to T cell engagers

In this study, we provide evidence that the catch bond phenomenon is not confined to membrane-bound TCRs. Notably, we demonstrate for the first time that a TCR-based BiTE can establish catch bonds with pMHC when tethered to CD3 on T cells. BiTEs engineered to exploit catch bond mechanics exhibited prolonged bond lifetimes, increased cytokine secretion, enhanced cytotoxic activity, and superior antitumor efficacy *in vivo* relative to affinity-matured BiTE counterparts. These findings broaden the therapeutic applicability of catch bond engineering beyond cellular therapies to include off-the-shelf, protein-based modalities, thereby providing a cost-effective and administratively versatile alternative. Furthermore, the efficacy of catch-bond-optimized BiTEs highlights the generalizability of the underlying mechanism, demonstrating that force-dependent enhancement of molecular interactions can be achieved across a variety of molecular architectures and valencies.

In conclusion, our study reveals that catch bonds play a critical role in modulating TCR sensitivity, and that engineering these catch bonds surpasses conventional affinity maturation approaches in enhancing TCR-driven antitumor efficacy. By redirecting the engineering paradigm from static affinity optimization to the dynamic stabilization of bonds under mechanical force, this approach aligns therapeutic development with the intrinsic mechanobiological principles governing TCR activation. This methodology is applicable to both cellular therapies, such as TCR-T, and bispecific engager molecules, thereby providing a versatile and rational framework for the advancement of next-generation TCR-based immunotherapies. As the field progresses toward more precise and safer immunotherapeutic strategies, catch bond engineering emerges as a promising and innovative avenue for future development.

## Methods

### Cell lines

The SKW3 cell line was obtained from DSMZ. The 293T, K562, NCI-H1299, SK-MEL-28, LN-18, A375, and Genti cell lines were purchased from the American Type Culture Collection (ATCC). The LentiX cell line was obtained from TaKaRa. Jurkat cells were kindly provided by the Chenqi Xu laboratory. Expi293F cells were purchased from Thermo Fisher Scientific.

SKW3, Jurkat, HCT116, K562, and T2 cells were cultured in complete RPMI 1640 medium (Gibco, Cat# C11875500BT) supplemented with 10% fetal bovine serum (FBS; Excell, Cat# FSD500) and penicillin/streptomycin (BBI, Cat# E607011-0100). LentiX, A375, and 293T cells were maintained in complete DMEM (Gibco, Cat# C11995500BT) supplemented with 10% FBS and penicillin/streptomycin. Expi293F and Genti cells were cultured in 293F Hi-Expression Medium (OPM Biosciences, Cat# AC601501).

### Animals

6- to 8-week-old female NOD.Cg-PrkdcscidIL 2rgtm1Sug/JicCrl (NOG) mice (#408) (Beijing VitalRiver Laboratory Animal Technology) were used for *in vivo* study. Animals were maintained in the animal facility of Center for Excellence in Molecular Cell Science, Chinese Academy of Sciences. All study protocols were approved by the Institutional Animal Care and Use Committee (IACUC) of Center for Excellence in Molecular Cell Science, Chinese Academy of Sciences.

### Antibodies

The following antibodies were used: anti-digoxigenin (Abcam, cat# ab420); from Biolegend: APC anti-human IgG Fc (cat# 410711), APC anti-human CD69 (cat# 310910), PE anti-human CD69 (cat# 310906), FITC anti-HA.11 epitope tag (clone 16B12, cat# 901507), APC anti-mouse TCR β chain (clone H57-597, cat# 109212), FITC anti-mouse TCR β chain (cat# 159706), PE anti-human CD107a (LAMP-1, cat# 328608), PE anti-human CD45 (cat# 304039), Brilliant Violet 605 anti-mouse IFN-γ (cat# 506542), PE/Cyanine7 anti-human TNF-α (cat# 502930), PE anti-human CD137 (4-1BB, cat# 309804), APC anti-human CD279 (PD-1, cat# 329908), PE anti-human α/β T cell receptor (cat# 306708), Brilliant Violet 421 anti-human TCR α/β (cat# 306722), APC anti-human TCR α/β (cat# 306718), anti-human CD3 (cat# 317326), anti-human CD28 (cat# 302934), Alexa Fluor 700 anti-human CD279 (PD-1, cat# 329951), Brilliant Violet 421 anti-human CD69 (cat# 310930), and PE anti-mouse TCR β chain (clone H57-597, cat# 109207); from Invitrogen: PE anti-mouse TCR β chain (cat# 12-5961-83), APC anti-mouse TCR β chain (cat# 17-5961-82), and streptavidin, R-phycoerythrin conjugate (SAPE, cat# S866); from BD Biosciences: BV421 anti-human CD8 (cat# 562428; also listed as BD cat# 562428); from Cell Signaling Technology: Alexa Fluor 647 TIM-3 (clone D5D5R, cat# 78226S); from eBioscience: Super Bright 600 anti-human CD366 (TIM3, clone F38-2E2, cat# 63-3109-42); and from SAILYBIO: APC anti-(G4S)n linker (cat# XY-Ab001-25T-APC).

### Establishment of TCR-transduced SKW3 cells

6x10^5^ lenti-X cells were seeded in 6-well plate with 2 mL completed DMEM medium. Next day, 750 ng pHR-TCR, 500 ng psPAX (Addgene, Cat# 12260), and 250 ng pMD2G (Addgene, Cat# 12259) were mixed with 1.5 μL Polyetherimide (PEI 1 mg/mL, Kyfora Bio, Cat# 24765-100) and 100 μL Opti-MEM (Gibco, Cat# 31985070). The mixture was vortexed and rested at room temperature for 20 min. The supernatant of lenti-X cell culture was replaced with 2 mL fresh completed RPMI medium. The DNA/PEI/Opti-MEM mixture was added into the lenti-X culture drop wisely. The lenti-X cells were cocultured at 37 ℃, 5% CO_2_ for 48 hours. The lentivirus supernatant was collected and filtered through 0.45 μm strainer to remove cell debris. 4 mL lentivirus supernatant were added into 10^6^ SKW3 T cells and cultured at 37 ℃, 5% CO_2_ for 48 hours. The expression of TCR were detected by anti-TCR staining and analyzed by flow cytometry (Thermo Fisher Attune).

### TCR-SKW3 activation experiments by peptide titration

Antigen-presenting cells (APCs) used in this paper were T2 or A375. APCs were pulsed with titrated peptides (Genscript) and cultured at 37 ℃, 5% CO_2_ for 3 hours. APCs were washed with cRPMI once to remove excess peptides. For CD69 measurement, 10^5^ APCs were cocultured with 10^5^ TCR-SKW3 T cells at 37 ℃, 5% CO_2_ for 14 hours. Cells were stained with anti-CD69 (Biolegend, Cat# 310910) and analyzed by flow cytometry.

### Establishment of primary TCR-T cells

TCR-T cells were produced in the lenti-X cells by generating lentiviral supernatant. In brief, 80% confluent lenti-X 10 cm dish was co-transfected with 10 μg pHR vector plasmid, 5 μg pVSVG (Addgene, Cat# 138479), and 7.5 μg pCMV-dR8.91 (Addgene, Cat# 8455) packaging plasmid DNA using Lipofectamine™ 3000 Reagent (Invitrogen, Cat# L3000008) according to manufacturer instructions. The 48- and 72-h viral supernatants were collected, centrifuged at 4000 rpm for 5 min, and filtered through a 0.45 μm PES membrane for either lentiviral infection of target cells or storage at -80°C. For viral infection, 5 million human PBMCs (SAILYBIO, Donor NO.W1274, IRB number: XF-WBC-220102) pre-stimulated with Dynabeads® Human T-Expander CD3/CD28 (Gibco, Cat# 11161D) at a ratio of 2.5:1 for 48 h were co-cultured with 2 mL completed X-VIVO 15 medium supplemented with 2 mM L-Glutamine (Gibco, Cat# A2916801), 1 mM sodium pyruvate (Beyotime, Cat# C0331-100ml), 1X NEAA (Gibco, Cat# 11140050), 55 μM 2-mercaptoethanol (Gibco, Cat# 21985023) and 100IU/mL IL-2 (Novoprotein, Cat# C013-10ug), along with 2mL viral supernatant containing 100IU/mL IL-2 and 8 μg/mL polybrene (Merck, Cat# TR-1003-G). The mixture was centrifuged at 800×g for 90 min at 30°C. On day 2 post-infection, TCR-T cell transduction efficiency was assessed by APC anti-mouse TCRβ (Biolegend, Cat# 109212).

### Primary TCR-T cells activation experiments *in vitro*

For intracellular cytokine staining, 5×10 MAGE-A4-specific TCR-T cells were co-cultured at a 1:1 ratio with A375 or T2 cells pre-treated with MAGEA4 peptide for 3 h in 96-well plates at 37°C for 6 h, supplemented with protein transport inhibitor (1:1000, BD GolgiPlug, Cat# 555029). After co-incubation, cells were surface-stained on ice with anti-CD8 BV421 (Biolegend, Cat# 562428) and anti-TCRβ APC. Subsequently, cells were treated with IC fixation buffer (Invitrogen, Cat# 00-8222-49) and permeabilization buffer (Invitrogen, Cat# 00-8333-56) sequentially, followed by intracellular staining with anti-TNFα PE-Cy7 (Biolegend, Cat# 502930). After staining, cells were analyzed by flow cytometry.

To assess the serial killing capacity of TCR-T cells, 5 × 10 TCR-T cells were co-cultured with A375 at the E: T ratio of 0:1, 0.5:1, 1:1, 2:1, and 4:1 in 96-well flat-bottom plates. After 24 h of incubation at 37°C, cells were collected, washed, and live T cells were counted. Fresh A375 was then added for a second round of challenge under the same conditions. This procedure was repeated for a total of 3 rounds. In each round, cytotoxicity was determined by Fixable Viability Dye eFluor™ 780 (Thermofisher, Cat# 65086518) by flow cytometry, and T cell phenotypes were detected by labeling anti-PD-1-AF700 (Biolegend, Cat# 329951) and anti-Tim-3-BV605 (Invitrogen, Cat# 406-3109-42).

### *In vivo* experiment of MAGE-A4-specific TCR-T cells

A375 cells (2×10^6^) were subcutaneously (s.c.) injected into the right flank of female NOG mice (6-8 weeks). After 7 days, the tumor-bearing mice were intravenously (i.v.) injected with PBS or 5×10^6^ MAGE-A4 TCR-T cells. Tumor volume was measured every 2-3 days using electronic calipers (formula: 0.5 × length (mm) × width (mm) × width (mm)). The experimental design was approved by IACUC (SIBCB-S650-2309-33).

### X-scan and off-target toxicity screening

First, a position-specific scoring matrix (PSSM) representing the amino acid preferences of the TCR-targeted peptide was generated using combinatorial peptide library screening, thereby defining residue preferences at each position of the peptide.

Second, the PSSM was used as input for proteome-wide screening using the HLA Compass platform. Briefly, the PSSM was uploaded to the “Toxicity Screening” module, the relevant HLA*02:01 allele was selected, and the scan was performed against the human proteome to identify candidate peptide sequences. These candidate peptides were subsequently scored using the python algorithm and ranked. The top-ranked peptide typically corresponds to the known endogenous target peptide (positive control), whereas peptides with scores approaching that of the target were considered high-confidence off-target candidates. Tissue expression profiles of these candidates (with green labels indicating expression in healthy tissues) were further examined to assess their potential biological relevance.

Third, candidate peptides with evidence of endogenous expression in human tissues were prioritized. The top 100 peptides with the highest off-target scores were synthesized and evaluated using peptide pulsing assays to assess TCR-T cell activation and potential off-target responses.

Finally, the experimental results were integrated with tissue expression data to assess overall off-target liability, with particular attention to whether high-scoring peptides are expressed in essential organs and whether the magnitude of T cell activation reaches functionally relevant thresholds.

### Alloreactivity screening

A panel of human tumor cell lines was used to assess the alloreactivity of Afamicel TCR-engineered T cells. The MAGE-A4 expression status and HLA-A*02:01 status of each cell line were confirmed prior to the assay. Target cells were pulsed with the indicated peptide at 10^-4 M and co-cultured with Afamicel T cells at an effector-to-target ratio of 1:1 for 12 h. T-cell activation was then evaluated by flow cytometric analysis of CD69 expression on mouse TCR Cβ-positive T cells.

### Protein expression and purification (Expi293/Expi293F GNTI^-^)

TCR4, TCR-5HTD, and TCR-5HIS were cloned into the pD649 backbone (Addgene, Cat# 156886). The pMHC was cloned as a single-chain trimer [peptide-(G₄S)₃-β₂m-(G₄S)₄-MHC extracellular domain (ECD)] into the pD649 backbone. Expi293/Expi293F GNTI⁻ cells were cultured in 293F Hi-exp medium at 37 °C under 5% CO₂ with agitation at 120 r.p.m. Transfection was performed when the cell density reached 2 × 10 cells/mL: 50 mL Opti-MEM, 1000 μg plasmids, and 3 mL PEI (1 μg/μL) were thoroughly mixed and incubated at room temperature for 20 min. The mixture was then added to 1 L of Expi293/Expi293F GNTI⁻ cell culture containing 2000 million cells. At 24 h post-transfection, 1000× enhancer 1 (200 mM valproic acid stock, Sparkjade, Cat# SJ-MN0055-500 mg) and 200× enhancer 2 (1.4 M sodium propionate stock, Adamas, Cat# 01592575) were added into the Expi293/Expi293F GNTI⁻ cell culture. The transfected cells were cultured for another 72 h before being collected. The cell culture supernatant was collected, and cell pellets were removed by centrifugation at 2000 g for 10 min at 4 ℃. To the supernatants of TCR4, TCR-5HTD, TCR-5HIS, and pMHC, 8 mL Ni-NTA was added, followed by incubation at 4℃ overnight. On the next day, the mixture was flowed through the column by gravity. The Ni-NTA resin was washed with 15 column volumes (CV) of HBS buffer (100 mM HEPES, 1.5 M NaCl, pH 7.2) containing 10 mM imidazole (Sangon, Cat# A600277-0500). Subsequently, the target protein was eluted with 6 CV of HBS buffer containing 300 mM imidazole. The eluted protein was concentrated and buffer-exchanged into HBS buffer using an Amicon concentrator. The protein concentration was finally measured on a Nanodrop. 1 μg of 3C protease was added to 50 μg of protein, mixed well, and incubated at room temperature for 2 h. The protein sample was added to pre-equilibrated Ni-NTA and stirred at room temperature for 30 min. Then, 1 CV of HBS buffer containing 10 mM imidazole was added, incubated for 10 min, and pushed out using a pressure pump; this step was repeated twice. The buffer was collected, concentrated to approximately 3 mL, and the protein concentration was measured on a Nanodrop. 1 μg of EndoH enzyme and 10× EndoH buffer were added to 100 μg of protein, mixed well, and incubated at room temperature for 1 h. The resulting protein sample was stored at 4 ℃. The protein was transferred to a 1.5 mL centrifuge tube with a filter membrane and centrifuged at 12,000 rpm for 10 min. The centrifuged protein was injected into a pre-equilibrated FPLC-S200 column using a 1 mL syringe, and the Elution program was run. The protein in the target centrifuge tubes was collected, concentrated to 10 mg/mL, and finally snap-frozen in liquid nitrogen before storage at -80 ℃.

### Surface plasma resonance

All SPR experiments were performed on a Biacore 8K at 25°C. A CM5 sensor chip was used throughout the study. The sensor chip surface was preconditioned with three consecutive injections of 50 mM NaOH containing 1 M NaCl for 30 seconds each at a flow rate of 100 µL/min. For ligand immobilization, the chip surface was activated by injecting a 1:1 mixture of 0.4 M EDC and 0.1 M NHS for 7 minutes at 10 µL/min. PMHC was diluted to 100 µg/mL in immobilization buffer and injected over the activated surface until the desired immobilization level 600 RU was achieved. Residual activated groups were blocked by injecting 1 M ethanolamine-HCl (pH 8.5) for 7 minutes. A reference surface was activated and ethanolamine-blocked without protein immobilization for background subtraction. For kinetic analysis, a series of concentrations of the TCR were prepared in running buffer HBS + 0.05% surfactant P20. The analyte was injected over the chip surface for an association phase of 120 seconds at a flow rate of 30 µL/min, followed by a dissociation phase of 600 seconds. Sensorgrams were processed by double referencing (subtraction of both reference surface and blank buffer injections). Kinetic rate constants (ka, kd) were determined by fitting the data globally to a 1:1 Langmuir binding model using Biacore Evaluation Software.

### Phospho-flow cytometry

Antigen-presenting cells (APCs) used in this paper were T2 or A375. APCs were pulsed with titrated peptides (Genscript) and cultured at 37 ℃, 5% CO_2_ for 3 hours. APCs were washed with cRPMI once to remove excess peptides. For pERK or pZAP70 measurement, 10^5^ APCs were cocultured with 10^5^ TCR-SKW3 T cells at 37 ℃, 5% CO_2_ for 15 min. Cells were immediately fixed by 4% PFA (Beyotime, Cat# P0099-100ml) and shaken for 15 min. The cells were washed with FACS washing buffer [PBS+0.5% BSA (Beyotime, Cat# ST023-200g)] and permeabilized with ice-cold methanol for 30 min on ice. The cells were washed with FACS washing buffer twice and stained with anti-pERK (clone 197G2, Cell Signaling Technology, Cat# 4377T) for 1 hour at room temperature with shaking. The cells were analyzed by flow cytometry (Thermo Fisher Attune).

### Magnetic Tweezer-Based Single-Molecule Experiments

The full plasmid constructs for TCR variants-HLA-A2-MAGE-A4 complexes were assembled sequentially as follows: His-tag and the TCRα variants at the N-terminus, followed by a biotinylated AviTag, the GS linker, HLA-A2-MAGE-A4, two tandem titin I27 domains as mechanical fingerprints, a SpyTag for covalent surface attachment. These sequences were cloned into pcDNA3.1 and co-expressed with the TCRβ variants. The recombinant proteins were collected and purified following similar protocol in Protein expression and purification parts.

A homemade magnetic tweezer (MT) setup, was employed for all single-molecule manipulation and measurement. System stability was enhanced by implementing an anti-drift procedure that utilizes piezo-driven focal plane adjustment, controlled via custom Labview software, allowing for stable force application over extended periods (days). In this configuration, the protein constructs were tethered within laminar flow channels between a Spy Catcher-coated cover glass and Neutravidin-coated superparamagnetic beads. A standard working buffer (1× PBS, 1% (m/v) BSA, 1 mM DTT, 10 mM L-ascorbic acid) was used for all experiments unless noted. Comprehensive methods regarding channel and bead preparation, sample handling, MT setup, force-jump experiments, and force calibration have been published previously^51^.

In order to measure the catch bond dynamics of TCR pMHC complexes, we quantified bond lifetimes using a force-jump assay. In each cycle, the complex was first allowed to form at low force (∼0.5 pN), followed by a rapid jump to a target force (5 pN, 10 pN, 15 pN, 20 pN), which was maintained until bond dissociation occurred. The dwell time prior to rupture was recorded as the bond lifetime, and repeated cycles enabled statistical analysis at each force.

### Small unilamellar vesicle (SUV) and supported lipid bilayer (SLB) preparation

To prepare small unilamellar vesicles (SUVs), phospholipids (90% POPC, 5% biotin-PE, and 5% DGS-NTA-Ni) were first dissolved in chloroform, followed by rotary evaporation to remove the solvent. The dried lipid film was then resuspended in PBS. The lipid suspension was sonicated for 60 min at 200 W and subsequently centrifuged at 20,000 × g at 4 °C for 2 h to remove large multilamellar vesicles (LMVs) and large unilamellar vesicles (LUVs). The supernatant containing SUVs was collected and stored at 4 °C under nitrogen protection.

For supported lipid bilayer (SLB) formation, glass-bottom 96-well plates were treated with isopropanol overnight and washed thoroughly with Milli-Q water. The plates were then incubated with 3 M NaOH for 1 h, followed by extensive washing with Milli-Q water. SUVs (13 nmol) were diluted in PBS and added to the wells for 25 min to allow SLB formation. Excess vesicles were removed by extensive washing with 1 mg/mL BSA in PBS (0.1% BSA/PBS).

### TIRF imaging of TCR clustering in antigen-free state

To analyze TCR clustering in the antigen-free state, Jurkat cells expressing different TCRs were seeded onto poly-L-lysine–coated glass-bottom dishes and fixed with 2× IC fixation buffer (Invitrogen, Cat# 00-8222) for 20 min. Cells were then stained overnight with anti-mouse TCRβ-PE (H57-597) in 1× permeabilization buffer (Invitrogen, Cat# 00-8333). After extensive washing with 0.1% BSA/PBS, cells were resuspended in PBS for imaging.

### TIRF imaging of immunological synapse

To analyze immunological synapse formation, Jurkat cells expressing CD2-BFP were lentivirally transduced with different TCR constructs. SLBs were blocked with 400 μL of 10 mg/mL BSA in PBS for 20 min at room temperature. After three washes with 0.1% BSA/PBS, streptavidin (2.5 ng/μL) was added and incubated for 20 min at room temperature. After washing, SLBs were incubated for 30 min at room temperature with 20 ng/μL biotinylated HLA-A2–MAGE-A4 pMHC (prepared in-house), 10 nM His-tagged ICAM-1, 10 nM His-tagged CD80, 10 nM His-tagged ICOSL, and 0.1 nM His-tagged CD58 (all from Sino Biological). Unbound proteins were removed by extensive washing with 0.1% BSA/PBS.

A total of 5 × 10^4^ Jurkat cells resuspended in DMEM were added onto the SLBs and incubated at 37 °C for 30 min to allow activation. Cells were then fixed with 2× IC fixation buffer for 20 min at room temperature and washed extensively to remove non-adherent cells. Cells were stained overnight with mouse anti-pCD3ζ (Y142)-Alexa Fluor 647 (BD, Cat# 558489) and rabbit anti-pZAP70 (Y493) (CST, Cat# 2704S) in 1× permeabilization buffer. After four washes with 0.1% BSA/PBS, anti-rabbit Cy3 secondary antibody was applied for 1 h at room temperature, followed by four additional washes.

### TIRF-SIM imaging and analysis

TIRF-SIM imaging was performed using a HIS-SIM system equipped with a 100× oil immersion objective. Pseudo-widefield images were generated by averaging 9 SIM raw frames and subsequently converted to 8-bit images. Image processing and analysis were carried out using ImageJ.

For quantification of TCR clustering in the antigen-free state, a threshold of 160 (gray value) was applied to the TCRβ channel to measure particle size. For immunological synapse size analysis, a threshold of 120 (gray value) was applied to the pCD3ζ channel. To quantify synaptic phosphorylation intensity, a circular region of interest (ROI) with a diameter of 6 μm was manually defined over the synapse area.

### Sortase assay

eSrtA was conjugated to cholesterol-PEG-NHS by incubation at equal mass in PBS with gentle stirring at 4 °C for 4 h. The reaction mixture was washed with PBS by ultrafiltration, supplemented with 5% glycerol, then aliquoted and stored at −80 °C. For cell-based assays, cells were incubated with cholesterol-conjugated protein at 37 °C for 15 min (in DMEM full medium), followed by washes 3 times with PBS before downstream experiments.

For SrtA-mediated proximity labeling, Streptococcus suis SrtA with S109A and Y130L mutations was used. Cell surface expression was achieved by direct cholesterol conjugation of purified eSrtA. For A375-TCR T cell labeling, TCR-T cells were co-cultured with cholesterol-conjugated-SrtA A375 cells (1:3). Biotin-AALPETGG (800 μM) was added, incubated for 2 h, then washed and analyzed by flow cytometry.

### Structural prediction

AlphaFold3 (https://alphafoldserver.com/) was used for the structure prediction of WT, TCR4 and 5mut in complex with HLA-A2-MAGE-A4^52^. Inputs included TCR α/β chains, peptide and MHC sequences. The top models were analyzed for interface residues using PyMOL.

### Protein Modeling and Molecular Dynamics Simulation

In this study, the TCR-pMHC simulation systems were constructed and analyzed. The wild-type structure was obtained as described below, while the corresponding mutated structures were generated by substituting the target residue with histidine, leucine, threonine, aspartic acid or glutamic acid.

For the WT complex, the coordinates of the TCR Vα and β domains (residues Q3-T113 in Vα and residues V2-E112 in Vβ), the HLA-A2 α1-2 domains (residues H3-Q180), and the antigen were taken from the complex structure predicted by the latest AlphaFold3 model^52^. After constructing the above complex structures, all TCR (Vαβ)-antigen-MHC (α1-2) systems were aligned so that the antigen axis, from the N-terminus to the C-terminus, was oriented along the +x direction. Molecular dynamics (MD) simulations were subsequently performed to refine the modeled structures and eliminate steric clashes or unrealistic interactions present in the initial complexes. All MD simulations were carried out using the AMBER 24 software package (University of California, San Francisco, USA) with the FF14SB force field^53^.

Each oriented complex was solvated in a rectangular TIP3P water box, maintaining a 10.0 Å buffer from the edges, and the system was further neutralized with NaCl. The system underwent three rounds of position-restrained energy minimization. Following this, the temperature was gradually increased to 300 K over 100 ps, which was followed by 10 ns of NVT equilibration and 10 ns of NPT equilibration. Subsequently, relaxation MD simulations were performed under conditions of 300 K and 1.0 atm. During this relaxation phase, bonds involving hydrogen atoms were constrained using the SHAKE algorithm^54^, enabling the use of a 2 fs integration time step. Non-bonded interactions were calculated with a 10.0 Å cutoff, while long-range electrostatic interactions were treated using the particle mesh Ewald (PME) method^55^. The Langevin thermostat was employed to maintain the temperature at 300 K, and the Berendsen barostat was applied to regulate the pressure at 1.0 atm. Based on the work of Sibener et al.^36^, a constant external force of 13.9 pN was applied to the TCR Vα and β chains in the +x direction during the relaxation phase. This force magnitude was insufficient to induce dissociation of the TCR-pMHC complex or cause significant displacement along the +x direction. All MD simulations were conducted on a high-performance computing cluster running the Linux operating system, with each system undergoing a total of 1500 ns of relaxation MD simulations.

The relaxed structures obtained from the relaxation simulations were subsequently used for shearing simulations. Prior to the shearing simulations, the relaxed structures were re-solvated in a cubic TIP3P water box with a side length of 120.0 Å, ensuring sufficient space for shearing in all directions. The final simulation box contained approximately 150,000 atoms. All shearing simulations were performed using steered molecular dynamics (SMD)^56^. In this study, constant-velocity SMD was employed to displace the center of mass of the TCR Vαβ relative to the MHC in the ±x and ±y directions.

The center of mass of the TCR Vαβ was connected to a point in space via a harmonic spring with a spring constant of 2.0 kcal/mol/Å2. This reference point was pulled at a constant velocity of 20 Å/ns in the specified direction. The SMD simulations were performed over a period of 2.5 ns, resulting in a total displacement of 50 Å in each direction.

Hydrogen bond analysis was conducted using the CPPTRAJ module^57^ in the AMBER 24 program. A hydrogen bond was defined as being formed if the distance between the donor and acceptor atoms was less than 3.2 Å, and the donor-hydrogen-acceptor angle was greater than 150°. A salt bridge is formed when the distance between oppositely charged residues is less than 3.2 Å. The interaction between TCR and pMHC was further characterized using MM-GBSA calculations^58^, with per-residue energy decomposition to determine key binding residues. The MM-GBSA calculations employed the GB model (GBOBC II) developed by Onufriev et al.^58^ for polar solvation energy, while the non-polar desolvation component was estimated based on the solvent-accessible surface area, using surften and surfoff parameters set to 0.005 and 0.0, respectively.

### Expression of bispecific T cell engager

A bispecific T cell engager (TCE) was generated by fusing the extracellular domain of the TCRα chain to an anti-CD3 scFv (clone UCHT1) via a (G4S)3 linker and cloning into the pD649 vector (pD649-TCEα), while the TCRβ chain extracellular domain was cloned into pD649 to generate pD649-TCEβ. To enhance stability and promote correct heterodimer assembly, multiple complementary strategies were combined: preservation of the native interchain disulfide bond (Cα94/Cβ130), incorporation of an additional commonly used stabilizing disulfide bond (T47C/S56C), and introduction of computationally derived stabilizing mutations (TRAC: S20F, T31I, A71T; TRBC: E17K, H22R, D38P, S53D) as previously described. Plasmids were co-transfected into Expi293F cells at a 1:1 ratio, and protein expression and purification were performed as described above.

### T cell activation experiments by bispecific T cell engager *in vitro*

For *in vitro* T cell activation assays, 5 × 10^4^ A375 cells were seeded into 96-well flat-bottom plates and allowed to adhere overnight. Primary human T cells were isolated from PBMCs by magnetic bead-based negative selection using the Human CD3^+^ T cell Isolation Kit (FiniGet) according to the manufacturer’s instructions, and 5 × 10^4^ T cells were added per well. Cells were co-cultured in the presence of indicated concentrations of TCE together with protein transport inhibitor (1:1000, BD GolgiPlug). After 18 h incubation, cells were first subjected to surface staining with anti-CD8 BV421 (Biolegend), followed by intracellular staining [anti-IFN-γ BV605 (Biolegend) and anti-TNF PE-Cy7 (Biolegend)] using Intracellular Flow Cytometry Fixation & Permeabilization Buffer Kit (Proteintech) according to the manufacturer’s instructions. Cells were analyzed by flow cytometry.

For *in vitro* T cell-mediated cytotoxicity assays, 5 × 10^4^ CFSE-labeled A375 cells were seeded into 96-well flat-bottom plates and allowed to adhere overnight. Primary human T cells isolated from PBMCs were then added at 5 × 10^4^ cells per well in the presence of indicated concentrations of TCE. After 42 h co-culture, cells were stained with 7-AAD and analyzed by flow cytometry. Viable tumor cells were defined as 7-AAD^-^ CFSE^+^ populations, and T cell-mediated cytotoxicity was calculated based on the reduction of viable target cells.

### Animal study of bispecific T cell engager

All animal experiments were performed in accordance with institutional guidelines and approved by the relevant animal care and use committee. Mice were maintained under specific pathogen–free conditions. Tumors were established by subcutaneous injection of 2 × 10^6^ A375 cells. On day 6, mice received 1 × 10^7^ human PBMCs via tail vein injection. TCE treatment was initiated on day 7 and administered intraperitoneally at 5 mg/kg every other day for a total of five doses. Tumor volume was measured every 2-3 days using electronic calipers (formula: 0.5 × length (mm) × width (mm) × width (mm)). Mice were monitored for general health and sacrificed when humane endpoints were reached. The experimental design was approved by IACUC (SIBCB-S650-2309-33).

## Acknowledgement

X.Z. is supported by the Strategic Priority Research Program of the Chinese Academy of Science (grant No. XDB0990000), National Natural Science Foundation of China (grant No. 82350107), the National Key Research and Development Projects of the Ministry of Science and Technology of the People’s Republic of China (grant No. 2024YFC2309200), Independent Project of Shanghai Sci-Tech Inno Center for Infection & Immunity ssIII-2024B01. W.H. is supported by the National Natural Science Foundation of China (92478124, 22473075) and Shanghai Pilot Program for Basic Research. This work is supported by the National Natural Science Foundation of China (grant no. 22573043, 22203089) to A.W. and the LiaoNing Revitalization Talents Program (No. XLYC2402023) to G. L.. B.S. is supported by the National Key Research and Development Program of China (2025YFC3409100) and the National Natural Science Foundation of China (32471508).

## Author contributions

X.Z. conceived the project. X.Z. wrote the manuscript with inputs from all the authors. T.H. performed histidine scanning, combinatory catch bond engineering, primary TCR-T cells in vitro and in vivo experiments, protein expression and purification, surface plasmon resonance experiment, phosphor-flow, X-scan, screening of off-target peptides, cell-cell interaction experiments. S.Wu. performed structure prediction and protein crystallization. M.F., X.W., L.Z., L.L., J.L., W.Y. performed isolation of tumor-infiltrating lymphocytes. Y.W. performed TCE expression and TCE in vitro and in vivo experiments. R.W., T.H., Y.B. performed catch bond measurements. D.Z., A.W. performed molecular dynamic simulations. Z.R. performed immunological synapse imaging. J.W. performed histidine scanning and X-scan. S.Wang. performed serine scanning, alloreactivity screening, protein purification. L.L. performed threonine scanning and protein purification. Y.Z. performed glutamine scanning, tyrosine scanning and protein purification. X.Z., W.H., A.W., C.X., G.L., F.S., B.S., S.L., H.Y. supervised the project. All authors edited the manuscript.

---

**Extended Data Fig.1.**
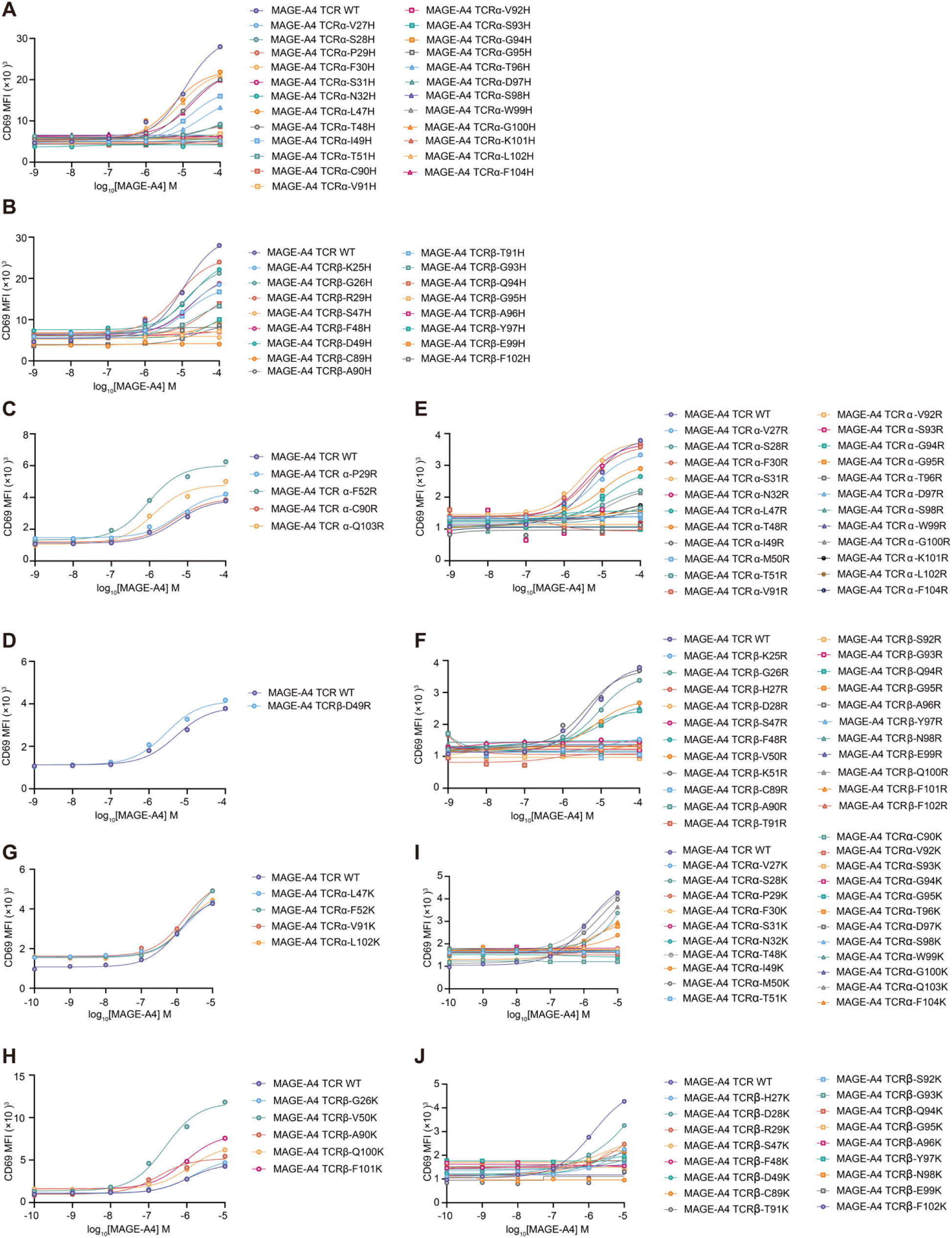
Histidine, arginine, and lysine scanning of TCR. (A-B) Histidine scanning of the WT TCR. Single-histidine-mutated WT TCR variants or the WT TCR were transduced into SKW3 cells. The SKW3 transfectants were activated by T2 cells pulsed with titrated MAGE-A4 peptides. The upregulation of CD69 in SKW3 transfectants was analysed by flow cytometry. The variants are classified based on their CD69 level: low on TCR α chain (A), low on TCR β chain (B). (C-F) Arginine scanning of the WT TCR. Single-arginine-mutated WT TCR variants or the WT TCR were transduced into SKW3 cells. The SKW3 transfectants were activated by T2 cells pulsed with titrated MAGE-A4 peptides. The upregulation of CD69 in SKW3 transfectants was analysed by flow cytometry. The variants are classified based on their CD69 level: high on TCR α chain (C), high on TCR β chain (D), low on TCR α chain (E), low on TCR β chain (F). (G-J) Lysine scanning of the WT TCR. Single-lysine-mutated WT TCR variants or the WT TCR were transduced into SKW3 cells. The SKW3 transfectants were activated by T2 cells pulsed with titrated MAGE-A4 peptides. The upregulation of CD69 in SKW3 transfectants was analysed by flow cytometry. The variants are classified based on their CD69 level: high on TCR α chain (G), high on TCR β chain (H), low on TCR α chain (I), low on TCR β chain (J). Data are representative of two experiments.

**Extended Data Fig.2.**
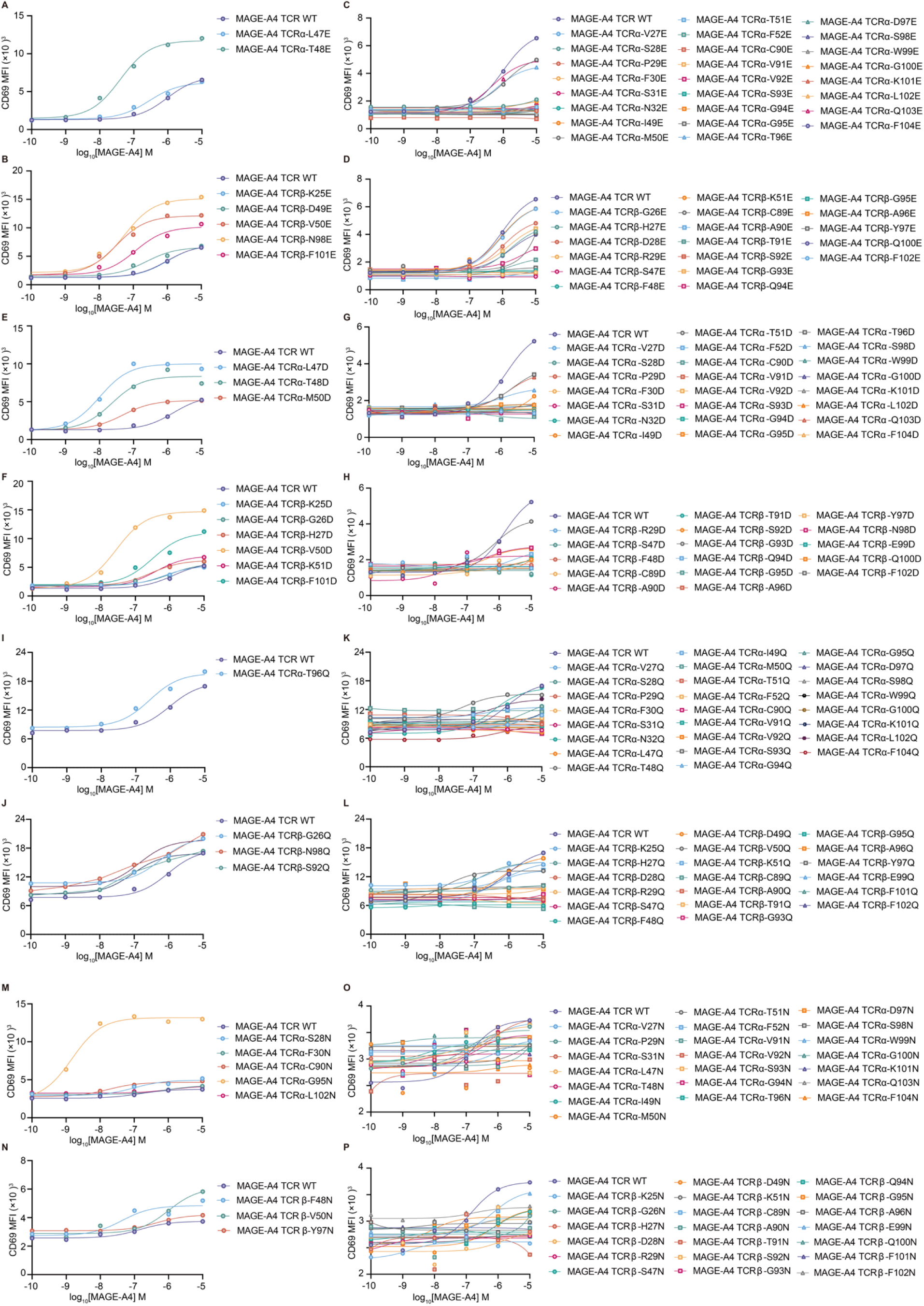
Glutamate, aspartate, glutamaine, and asparagine scanning of TCR. (A-D) Glutamate scanning of the WT TCR. Single-glutamate-mutated WT TCR variants or the WT TCR were transduced into SKW3 cells. The SKW3 transfectants were activated by T2 cells pulsed with titrated MAGE-A4 peptides. The upregulation of CD69 in SKW3 transfectants was analysed by flow cytometry. The variants are classified based on their CD69 level: high on TCR α chain (A), high on TCR β chain (B), low on TCR α chain (C), low on TCR β chain (D). (E-H) Aspartate scanning of the WT TCR. Single-aspartate-mutated WT TCR variants or the WT TCR were transduced into SKW3 cells. The SKW3 transfectants were activated by T2 cells pulsed with titrated MAGE-A4 peptides. The upregulation of CD69 in SKW3 transfectants was analysed by flow cytometry. The variants are classified based on their CD69 level: high on TCR α chain (E), high on TCR β chain (F), low on TCR α chain (G), low on TCR β chain (H). (I-L) Glutamine scanning of the WT TCR. Single-glutamine-mutated WT TCR variants or the WT TCR were transduced into SKW3 cells. The SKW3 transfectants were activated by T2 cells pulsed with titrated MAGE-A4 peptides. The upregulation of CD69 in SKW3 transfectants was analysed by flow cytometry. The variants are classified based on their CD69 level: high on TCR α chain (I), high on TCR β chain (J), low on TCR α chain (K), low on TCR β chain (L). (M-P) Asparagine scanning of the WT TCR. Single-asparagine-mutated WT TCR variants or the WT TCR were transduced into SKW3 cells. The SKW3 transfectants were activated by T2 cells pulsed with titrated MAGE-A4 peptides. The upregulation of CD69 in SKW3 transfectants was analysed by flow cytometry. The variants are classified based on their CD69 level: high on TCR α chain (G), high on TCR β chain (H), low on TCR α chain (I), low on TCR β chain (J). Data are representative of two experiments.

**Extended Data Fig.3.**
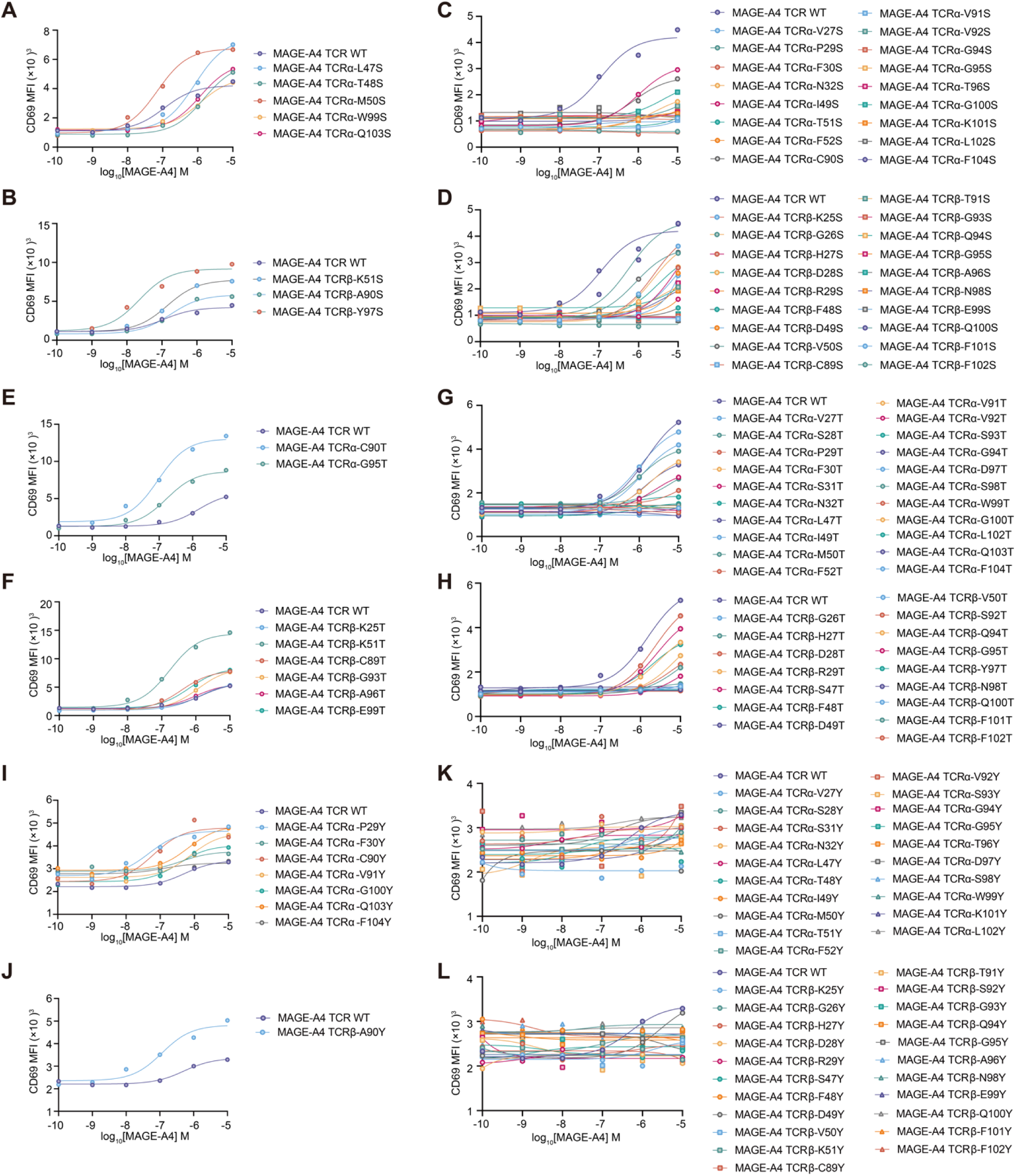
Serine, threonine, and tyrosine scanning of TCR. (A-D) Serine scanning of the WT TCR. Single-serine-mutated WT TCR variants or the WT TCR were transduced into SKW3 cells. The SKW3 transfectants were activated by T2 cells pulsed with titrated MAGE-A4 peptides. The upregulation of CD69 in SKW3 transfectants was analysed by flow cytometry. The variants are classified based on their CD69 level: high on TCR α chain (A), high on TCR β chain (B), low on TCR α chain (C), low on TCR β chain (D). (E-H) Threonine scanning of the WT TCR. Single-threonine-mutated WT TCR variants or the WT TCR were transduced into SKW3 cells. The SKW3 transfectants were activated by T2 cells pulsed with titrated MAGE-A4 peptides. The upregulation of CD69 in SKW3 transfectants was analysed by flow cytometry. The variants are classified based on their CD69 level: high on TCR α chain (E), high on TCR β chain (F), low on TCR α chain (G), low on TCR β chain (H). (I-L) Tyrosine scanning of the WT TCR. Single-tyrosine-mutated WT TCR variants or the WT TCR were transduced into SKW3 cells. The SKW3 transfectants were activated by T2 cells pulsed with titrated MAGE-A4 peptides. The upregulation of CD69 in SKW3 transfectants was analysed by flow cytometry. The variants are classified based on their CD69 level: high on TCR α chain (I), high on TCR β chain (J), low on TCR α chain (K), low on TCR β chain (L). Data are representative of two experiments.

**Extended Data Fig.4.**
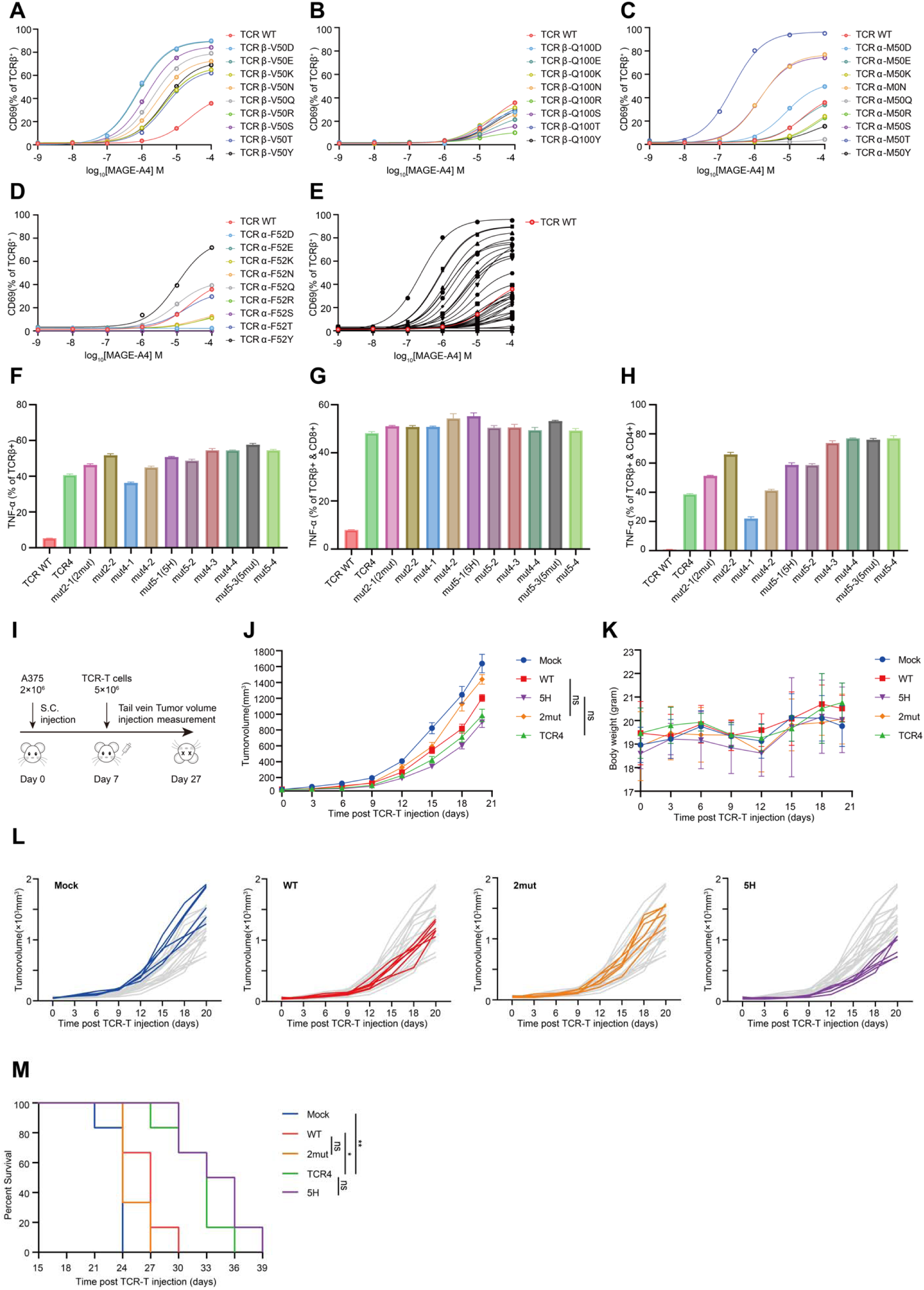
Combinatory catch bond engineering of TCR. (A) TCRβ-V50 residue was mutated to different polar or charged amino acids. Different TCR variants-transduced SKW3 cells were cocultured with T2 cells pulsed with titrated MAGE-A4 peptides. SKW3 cells were stained with anti-CD69-APC. (B) TCRβ-Q100 residue was mutated to different polar or charged amino acids. Different TCR variants-transduced SKW3 cells were cocultured with T2 cells pulsed with titrated MAGE-A4 peptides. SKW3 cells were stained with anti-CD69-APC. (C) TCRα-M50 residue was mutated to different polar or charged amino acids. Different TCR variants-transduced SKW3 cells were cocultured with T2 cells pulsed with titrated MAGE-A4 peptides. SKW3 cells were stained with anti-CD69-APC. (D) TCRα-F52 residue was mutated to different polar or charged amino acids. Different TCR variants-transduced SKW3 cells were cocultured with T2 cells pulsed with titrated MAGE-A4 peptides. SKW3 cells were stained with anti-CD69-APC.(E) Results in (A)-(D) were put together. (F) Several TCR variants with combinatory mutagenesis were transduced into primary T cells and cocultured with A375 cells. TNF expression in all T cells were stained and analyzed by flow cytometry. (G) Several TCR variants with combinatory mutagenesis were transduced into primary T cells and cocultured with A375 cells. TNF expression in CD8^+^ T cells were stained and analyzed by flow cytometry. (H) Several TCR variants with combinatory mutagenesis were transduced into primary T cells and cocultured with A375 cells. TNF expression in CD4^+^ T cells were stained and analyzed by flow cytometry. Data are representative of two experiments. (I) Schematic of testing primary TCR-T cell therapy in tumor-bearing immunodeficient mice *in vivo*. (J) Tumor volume of mice post TCR-T cells injection. Immunodeficient mice were implanted with A375 tumor cells followed by adoptive transfer of TCR variants-transduced primary T cells. (K) Body weight of mice post TCR-T cells injection. (L) Tumor volume of individual mouse post TCR-T cells injection. (M)Survival curve of mice post TCR-T cells injection. Data are representative of two experiments. Data are shown as means ± SD of technical duplicates. Statistical analysis was performed using two-tailed unpaired Student’s *t*-tests. n.s., not significant; \**P* < 0.05; \*\**P* < 0.01; \*\*\**P* < 0.001; \*\*\*\**P* < 0.0001.

**Extended Data Fig.5.**
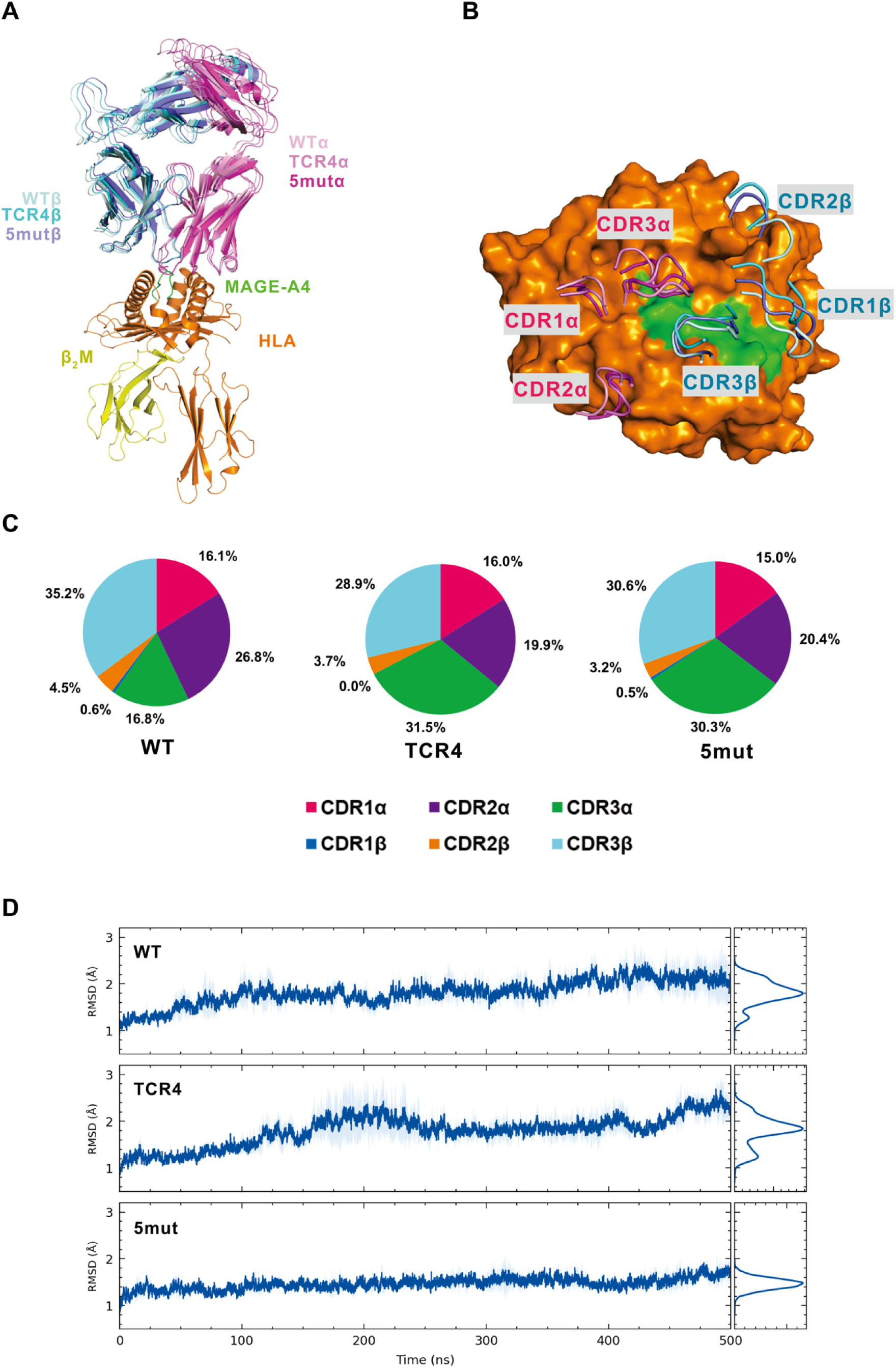
AlphaFold3-based structural models and MD stability analysis of TCR–pMHC complexes. (A) AlphaFold3-predicted structures of TCR WT and variants in complex with pMHC, superimposed on the HLA-A2 molecule. (B) Binding footprints of CDR loops from TCR WT and variants on the pMHC surface. Structural alignment shows high similarity (Cα RMSD: ∼1.0-1.4 Å), indicating minimal global conformational differences. Colors are the same as in (A). (C) Relative contributions of individual CDR loops to TCR-pMHC interactions. Mutations redistribute interaction contributions, particularly within CDR2α, CDR3α, and CDR3β. (D) Time evolution of Cα RMSD during 500 ns relaxation MD simulations. Each system represents the average of three independent trajectories, demonstrating stable conformational sampling across all complexes.

**Extended Data Fig.6.**
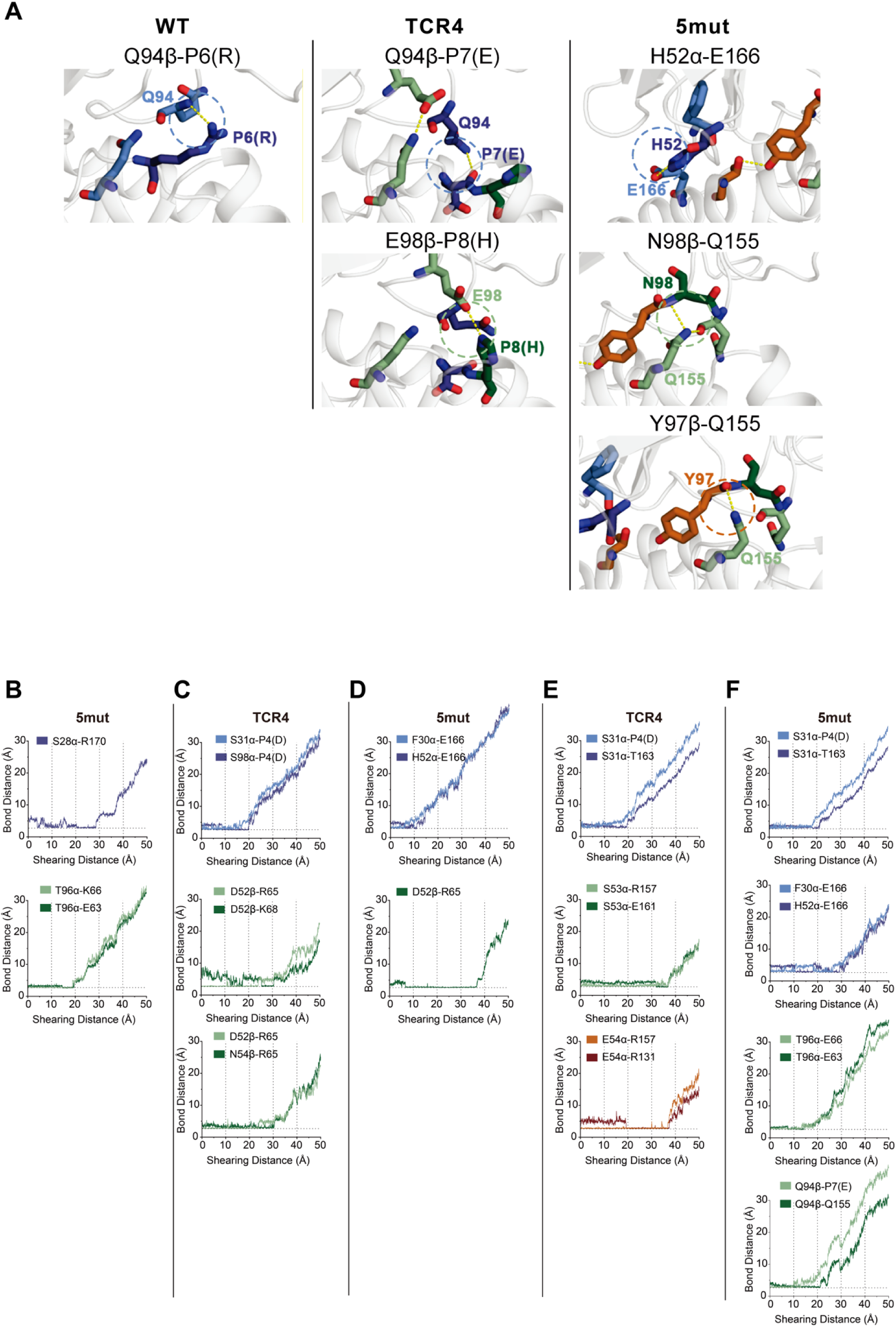
Representative snapshots and interaction dynamics during SMD simulations. (A) Representative snapshots illustrating key stages of TCR-pMHC dissociation under applied force, highlighting the rupture of original interactions and formation of new contacts (catch bonds). (B) -(F) Bond distance versus shearing distance plots for representative catch bonds under different shear directions: (B) 5mut under −x direction; (C) TCR4 under +y direction; (D) 5mut under +y direction; (E) TCR4 under −y direction; (F) 5mut under−y direction. Horizontal gray dashed lines indicate equilibrium bond distances. These plots demonstrate mutation-dependent formation of transient stabilizing interactions.

**Extended Data Fig.7.**
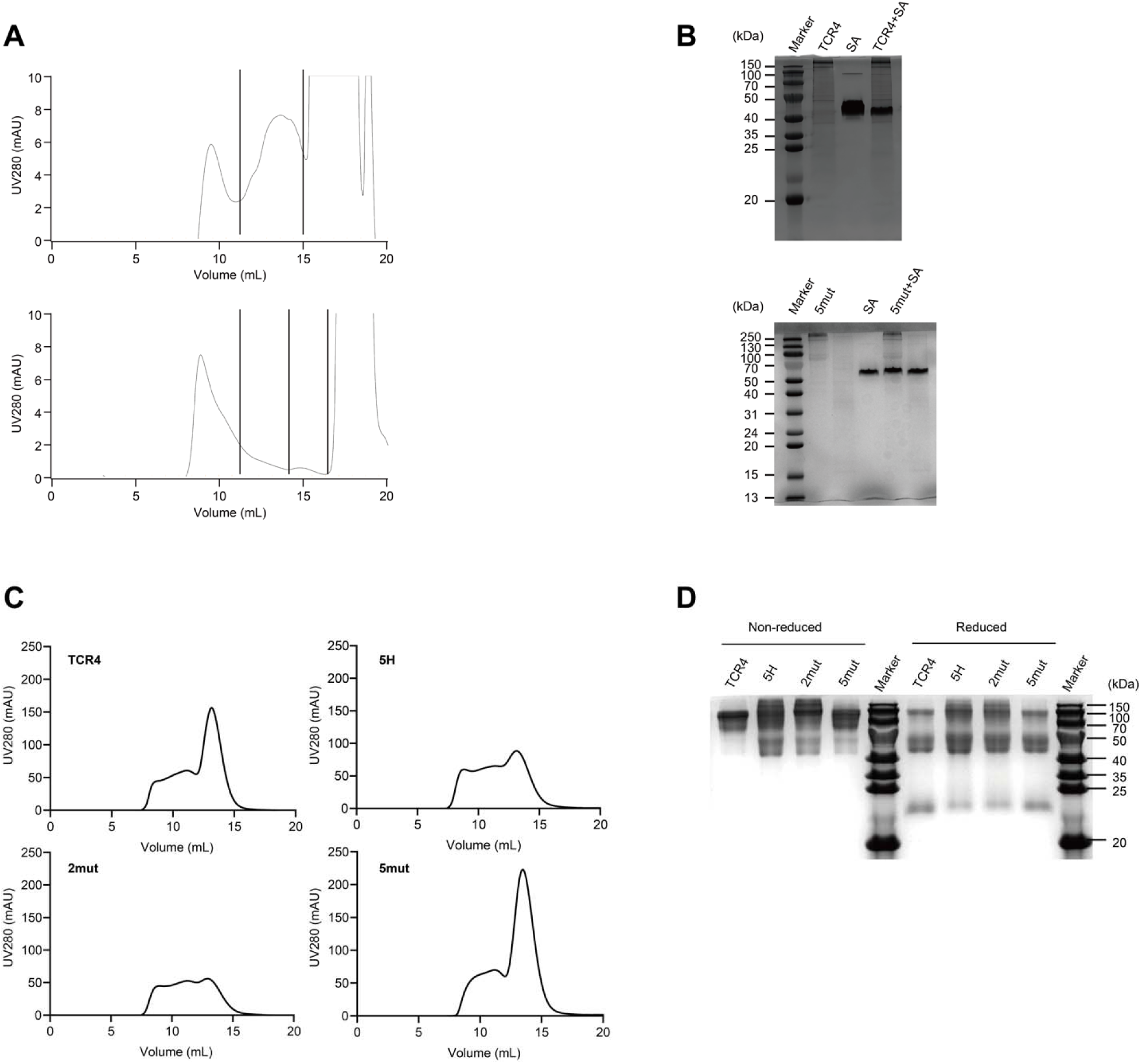
Expression of proteins for magnetic tweezers and expression of T cell engagers. (A) Purification of TCR for magnetic tweezers on fast-performance liquid chromatography. (B) Validation of TCR for magnetic tweezers expression by sodium dodecyl sulfate-polyacrylamide gel electrophoresis (SDS-PAGE). (C) Purification of T cell engagers on fast-performance liquid chromatography. (D) Validation of T cell engagers expression by sodium dodecyl sulfate-polyacrylamide gel electrophoresis (SDS-PAGE).

**Table S1.** Specific mutation sites and amino acid alterations in different MAGE-A4 TCR mutants.

|  |  |
| --- | --- |
| mut2-1 | TCRβ-V50H & TCRα-M50T |
| mut2-2 | TCRβ-V50D & TCRα-M50T |
| mut4-1 | TCRβ-V50H & TCRα-M50H , F52H , Q103H |
| mut4-2 | TCRβ-V50D & TCRα-M50H , F52H , Q103H |
| mut4-3 | TCRβ-V50H & TCRα-M50T , F52H , Q103H |
| mut4-4 | TCRβ-V50D & TCRα-M50T , F52H , Q103H |
| mut5-1 | TCRβ-V50H , Q100H & TCRα-M50H , F52H , Q103H |
| mut5-2 | TCRβ-V50H , F101H & TCRα-M50H , F52H , Q103H |
| mut5-3 | TCRβ-V50D , Q100H & TCRα-M50T , F52H , Q103H |
| mut5-4 | TCRβ-V50D , F101H & TCRα-M50T , F52H , Q103H |
| mut6 | TCRβ-V50H , Q100H , F101H & TCRα-M50H , F52H , Q103H |
| TCR4 | TCRβ-N98E & TCRα-M50L |

**Table S2.**
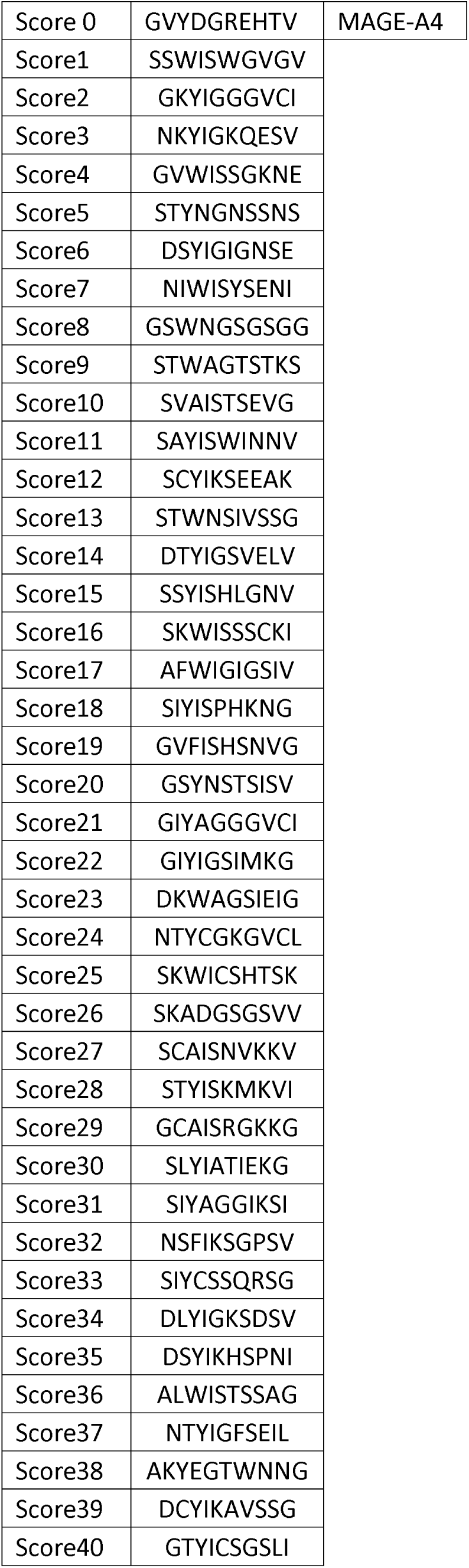
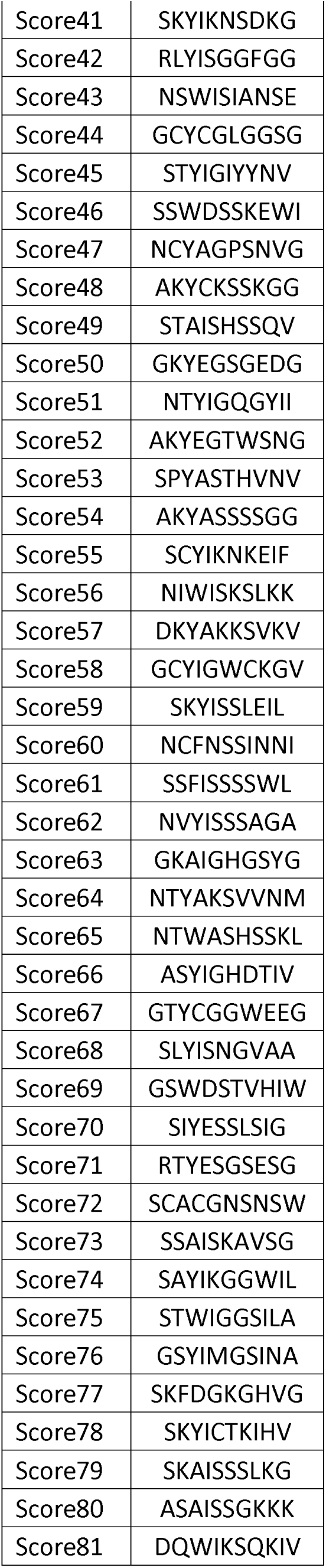
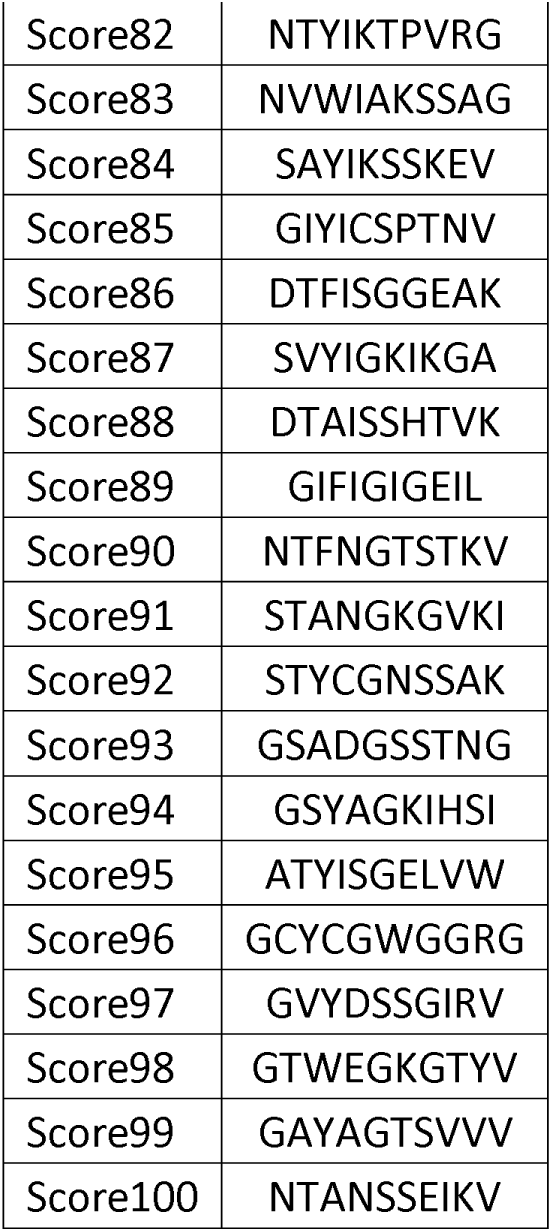
Ranked predicted off-targeted peptides. A position-specific scoring matrix (PSSM) derived from X-scan data was used to score all possible peptides in the human proteome. Peptides were encoded using one-hot representation and scored by matrix multiplication with the PSSM. The top 100 peptides with the highest predicted scores were selected as candidate off-targets for downstream analysis.

**Table S3.** Interfacial hydrogen bonds and energetic contributions from relaxation MD simulations. Hydrogen bonds at the TCR-pMHC interface identified from MD simulations for WT TCR, TCR4, and 5mut. For each interaction, the occupancy (frequency) and estimated energetic contribution (kcal/mol) are reported. Total energetic contributions highlight enhanced binding in TCR4 (∼−35 kcal/mol) and intermediate stabilization in 5mut (∼−25 kcal/mol), compared to WT (∼−17 kcal/mol).

| Hydrogen Bonds |  | TCR WT |  | TCR 4 |  | TCR 5mut |  |
| --- | --- | --- | --- | --- | --- | --- | --- |
|  |  | Frequency | Contribution<br>(kcal/mol) | Frequency | Contribution<br>(kcal/mol) | Frequency | Contribution<br>(kcal/mol) |
| F30α | E166 | 23.88% | -1.43 | 10.18% | -0.35 | 33.52% | -1.22 |
| D97α | K66 | 0.00% | 0.44 | 91.89% | -6.59 | 0.00% | 2.06 |
| Q94β | Q155 | 47.75% | -4.73 | 15.26% | -3.67 | 60.46% | -6.25 |
| Y97β | P4 (T) | 11.25% | -7.94 | 99.66% | -15.66 | 99.96% | -16.70 |
| Y97β | T163 | 67.41% | -9.81 | 88.51% | -13.45 | 97.63% | -14.89 |
| N/E98β | Q155 | 14.88% | -2.68 | 33.29% | -3.99 | 7.30% | -1.81 |
| N/E98β | P6 (R) | 1.85% | -0.59 | 55.53% | -5.45 | 0.69% | 0.18 |
| All Contributions<br>(kcal/mol) |  |  | -17.09 |  | -35.00 |  | -25.45 |

**Table S4.** Catch bond interactions formed during SMD simulations. Summary of catch bonds formed during forced dissociation of TCR variants from pMHC under shear in the ±x and ±y directions. For each system, the number and identity of newly formed interactions are listed, revealing mutation-dependent differences in dynamic interaction remodeling.

| Direction | System | WT/Mut | No. of Catch Bond | Catch Bond Details |  |  |  |
| --- | --- | --- | --- | --- | --- | --- | --- |
| +x | sys1 | TCR WT | 1 | Q94β-P6 (R) | | | |
| | sys2 | TCR 4 | 2 | Q94β-P7 (E) | E98β-P8 (H) | | |
| | sys3 | TCR 5mut | 3 | H52α-E166 | Y97β-Q155 | N98β-Q155 | |
| -x | sys1 | TCR WT | 0 |  |  |  |  |
|  | sys2 | TCR 4 | 0 |  |  |  |  |
| | sys3 | TCR 5mut | 2 | S28α-R170 | T96α-E63 | | |
| +y | sys1 | WT | 0 |  |  |  |  |
| | sys2 | TCR 4 | 3 | S98α-P4 (D) | D52β-K68 | N54β-R65 | |
| | sys3 | TCR 5mut | 2 | H52α-E166 | D52β-R65 | | |
| -y | sys1 | TCR WT | 0 |  |  |  |  |
| | sys2 | TCR 4 | 3 | S31α-T163 | S53α-E161 | E54α-R131 | |
| | sys3 | TCR 5mut | 4 | S31α-T163 | H52α-E166 | T96α-E63 | Q94β-Q155 |

**Table S5.** Quantification of catch bond formation under directional shear. Summary of the number of catch bonds formed by each TCR variant under different shear directions (±x, ±y). The 5mut variant exhibits the highest average number of catch bonds, consistent with its enhanced ability to sustain interactions under force.

| System | WT/Mut | +x | -x | +y | -y | Average |
| --- | --- | --- | --- | --- | --- | --- |
| sys1 | TCRWT | 1 | 0 | 0 | 0 | 0.25 |
| sys2 | TCR4 | 2 | 0 | 3 | 3 | 2.00 |
| sys3 | TCR 5mut | 3 | 2 | 2 | 4 | 2.75 |

**Movie S1.**
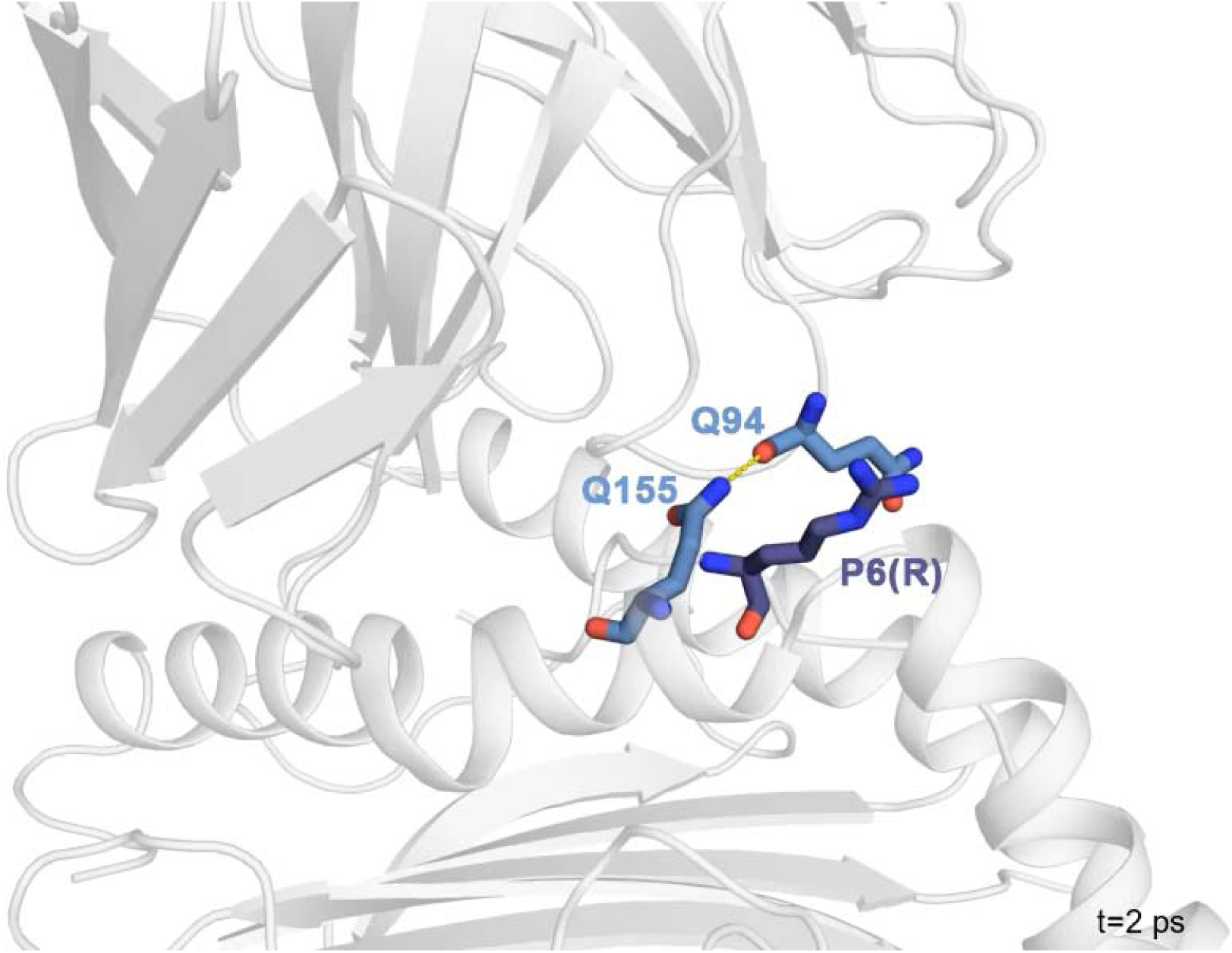
Molecular dynamics simulation of WT TCR binding to HLA-A2-MAGE-A4 under force. The shearing of WT TCR (top, gray) along the N-term to C-term antigen axis (central, gray) with respect to HLA-A2 (bottom, gray) leads to several molecular catch bonds for MAGE-A4. In hotspot residues critical for catch bond formation, colored in purple, one residue in the starting bond (lighter hue) breaks its interaction and forms a new interaction with a bonding partner (darker hue). Yellow dashed lines connect the forming bonds at their equilibrium distances. The catch bonds between WT TCR and HLA-A2-MAGE-A4 exhibit transient duration, failing to augment the binding stability of the complex.

**Movie S2.**
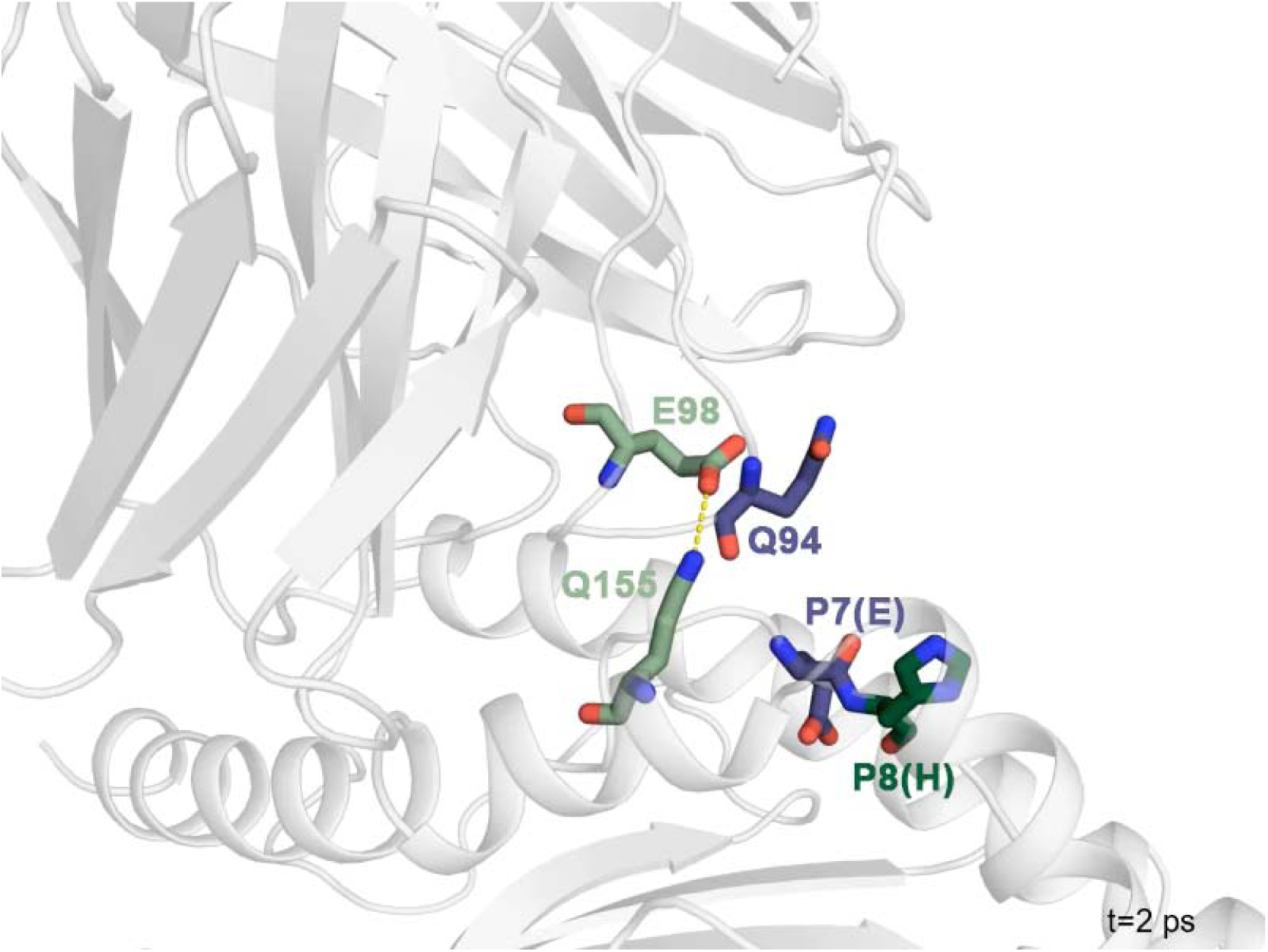
Molecular dynamics simulation of TCR4 Binding to HLA-A2-MAGE-A4 under force. The shearing of TCR4 (top, gray) along the N-term to C-term antigen axis (central, gray) with respect to HLA-A2 (bottom, gray) leads to two molecular catch bonds for MAGE-A4. In hotspot residues critical for catch bond formation, colored in green and purple, one residue in the starting bond (lighter hue) breaks its interaction and forms a new interaction with a bonding partner (darker hue). Yellow dashed lines connect the forming bonds at their equilibrium distances. The catch bonds between TCR4 and HLA-A2-MAGE-A4 demonstrate prolonged durability and increased frequency, providing sustained resistance against relative shearing.

**Movie S3.**
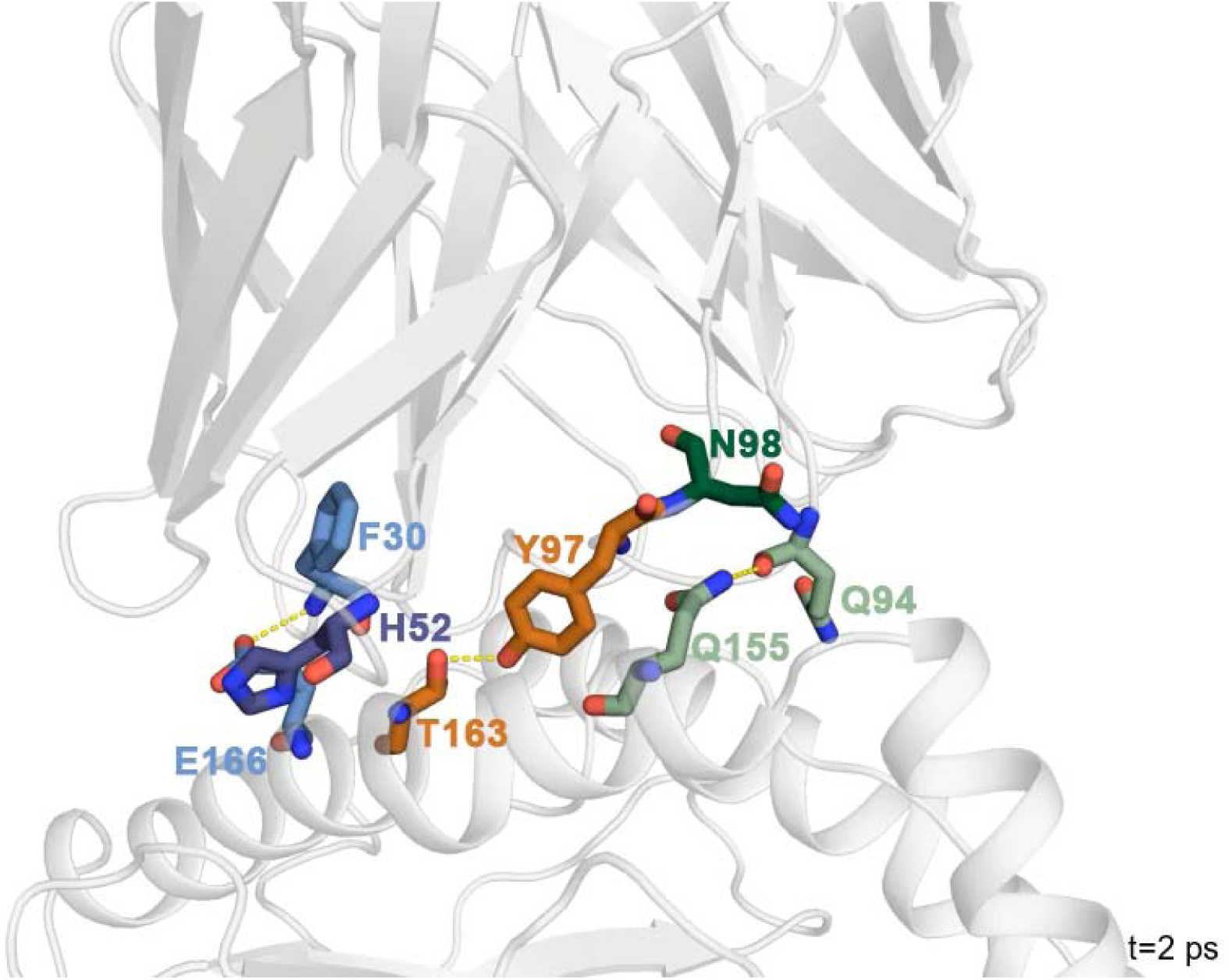
Molecular dynamics simulation of 5mut binding to HLA-A2-MAGE-A4 under force. The shearing of 5mut (top, gray) along the N-term to C-term antigen axis (central, gray) with respect to HLA-A2 (bottom, gray) leads to three molecular catch bonds for MAGE-A4. In hotspot residues critical for catch bond formation, colored in orange, green and purple, one residue in the starting bond (lighter hue) breaks its interaction and forms a new interaction with a bonding partner (darker hue). Yellow dashed lines connect the forming bonds at their equilibrium distances. The catch bonds between 5mut and HLA-A2-MAGE-A4 demonstrate prolonged durability and increased frequency, providing sustained resistance against relative shearing.

